# Within-host viral growth and immune response rates predict FMDV transmission dynamics for African Buffalo

**DOI:** 10.1101/2022.12.02.518883

**Authors:** Joshua C. Macdonald, Hayriye Gulbudak, Brianna Beechler, Erin E. Gorsich, Simon Gubbins, Eva Pérez-Martin, Anna E. Jolles

**Author notes:** Corresponding author; HG, AJ.

## Abstract

Infectious disease dynamics operate across biological scales: pathogens replicate within hosts but transmit among populations. Functional changes in the pathogen-host interaction thus generate cascading effects across organizational scales. We investigated within-host dynamics and among-host transmission of three strains (SAT-1, 2, 3) of foot-and-mouth disease viruses (FMDVs) in their wildlife host, African buffalo. We combined data on viral dynamics and host immune responses with mathematical models to ask (i) How do viral and immune dynamics vary among strains?; (ii) Which viral and immune parameters determine viral fitness within hosts?; and (iii) How do within-host dynamics relate to virus transmission? Our data reveal contrasting within-host dynamics among viral strains, with SAT-2 eliciting more rapid and effective immune responses than SAT-1 and SAT-3. Within-host viral fitness was overwhelmingly determined by variation among hosts in immune response activation rates but not by variation among individual hosts in viral growth rate. Our analyses investigating across-scale linkages indicate that viral replication rate in the host correlates with transmission rates among buffalo and that adaptive immune activation rate determines the infectious period. These parameters define the virus’s relative basic reproductive number (ℛ_0_), suggesting that viral invasion potential may be predictable from within-host dynamics.

## Introduction

Linking pathogen dynamics across biological scales, from cellular and molecular interactions within the host’s tissues to transmission among individuals and populations, is critical to understanding ecological and evolutionary trajectories of host-pathogen systems and represents a central challenge in disease ecology (Plowright et al., 2008; Pongsiri et al., 2009; Wale and Duffy, 2021). Multi-scale models of infectious disease dynamics seek to address this challenge by linking mechanistic models representing pathogen-host interactions at cellular to population scales. Developing the mathematical tools for connecting dynamic processes operating at vastly different temporal and spatial scales has been an active focus in infectious disease modeling (Agyingi et al., 2020; Browne and Cheng, 2020; Garabed et al., 2020; Garira, 2020; Jia et al., 2020; Kadelka and M Ciupe, 2019; Rivera et al., 2020; Versypt, 2021; Xue and Xiao, 2020). However, these theoretical innovations have yet to be matched by empirical data generation, providing integrated data sets that consistently document infection processes in the same host-pathogen system across organizational scales.

In this study, we leveraged experimental data on within-host dynamics and among-host transmission of three strains of foot-and-mouth disease viruses (FMDVs) in their wild reservoir host, African buffalo (*Syncerus caffer*). We constructed a data-driven mathematical model to understand the interplay between viral population growth and its limitation by the host’s immune responses. We then investigated to what extent parameters capturing within-host viral dynamics can predict variation in viral fitness at within- and among-host scales. Within the host, we define *fitness* as viral production in terms of peak and cumulative viral load. As such, these are both measures of relative success between viral strains. At the population scale, we assessed fitness in terms of the relative basic reproductive number ℛ_0_, assuming constant contact rates and population densities (note, as with any proxy, ℛ_0_ is an imperfect measure of fitness, see Lion and Metz, 2018, for a discussion of its limitations alternative metrics). The basic reproductive number is defined as the mean number of secondary infections caused by a single infected host in a wholly susceptible population, ℛ_0_ represents the pathogen’s ability to invade susceptible host populations (see Diekmann et al., 1990). Thus, our approach connects within-host viral dynamics to the potential for pathogen spread in host populations, providing a first step towards integrating data and disease dynamic models across biological scales in this study system.

FMDVs in African buffalo provide a tractable model system for studying natural populations’ multi-scale infection processes. FMDVs are highly contagious viruses that cause clinical disease and substantial production losses in domestic ungulates, while endemic infections in their wildlife reservoir tend to be milder (Coetzer et al., 1994; Gainaru et al., 1986). FMDVs are ubiquitous in African buffalo (Coetzer et al., 1994), with three distinct serotypes circulating in wild buffalo populations essentially independently in Southern Africa (Maree et al., 2016; Thomson et al., 1992; Vosloo et al., 1996), allowing meaningful comparisons across sympatric viral strains. FMD is the most important trade-restricting livestock disease globally. As a result, well-established methods exist for virus culture, experimental challenges, diagnostics, and quantifying immune responses (Couch et al., 2017; Glidden et al., 2018; Jolles et al., 2021; Maree et al., 2016; Stenfeldt et al., 2011).

Previous work has shown that FMDV strains vary substantially in their transmission dynamics among buffalo hosts (Jolles et al., 2021), and viral proliferation and immune response patterns in buffalo have been described (Perez-Martin et al., 2022). However, the functional interplay of within-host viral and immune dynamics has yet to be evaluated in buffalo, compared among Southern African Territories (SAT) serotypes, or aligned with population-level disease dynamics. In this study, we combined experimental infection data and a mechanistic mathematical model to ask: (i) How do viral and immune dynamic interactions vary among FMDV strains? (ii) Which viral and immune parameters determine viral fitness within hosts?; and (iii) How do within-host dynamics relate to virus transmission among hosts? Our data and models show that variation among viral strains in dynamics within buffalo hosts is reflected in variation in transmission dynamics among hosts, demonstrating agreement in viral dynamics across biological scales.

## Materials and methods

### The data

We conducted transmission experiments in which time series data were collected to quantify viral and immune kinetics within each host and to estimate epidemiological parameters such as transmission rate and infectious period for one strain of each serotype for primary (acute) FMDV infection. By acute infection, we mean the time from the beginning of infection to the clearance of the virus from the blood of each host. Population scale data and parameter estimates were published along with our prior work (Jolles et al., 2021). Detailed experimental protocols are available in published form (see Jolles et al., 2021; Perez-Martin et al., 2022); we summarize the most pertinent details here. See the SI of this work for a detailed explanation of the computational methods.

Four buffalo were needle-infected for each strain and allowed to contact four naive buffalo. These naive buffalo were then captured on contact days 0, 2, 4, 6, 9, and 12 during the acute phase and again at 28 days post-contact to measure viral and immune parameters. While several markers for the immune responses were considered (Macdonald et al., 2024; Perez-Martin et al., 2022), here we focus on the innate response as measured by Haptoglobin [log_10_ *μg*/*mL*], virus [log_10_(genome copies/mL)], and adaptive response as measured by virus neutralization titer [log_10_(VNT)]. Fever periods were estimated based upon continuously gathered temperature data via surgically implanted temperature loggers in each buffalo (see Perez-Martin et al., 2022, for details). The impact of the route of infection on model parameters is discussed extensively in the supplementary information. We find significant differences between the needle- and contact-infected hosts and focus our analysis here on the contact-infected hosts (i.e., animals infected via a natural route of infection).

### The mathematical model

Within-host tissue tropism in FMDV is complex (but see Li et al., 2021). Generally, within a given host, FMDVs first target pharyngeal epithelium and, then, for primary infections, spread to many other cell types. Hosts have “acute infection,” i.e., are viremic in both domestic cattle and buffalo for 6 to 12 days (including incubation) (Perez-Martin et al., 2022; Yadav et al., 2019). Our model (see system of equations (1), Tab. 1 and Fig. 1) includes the pathogen population [*P*(*τ*)], innate [*I*(*τ*)], and adaptive [*A*(*τ*)] immune effectors, where *τ* refers to time since infection of the host, and initial conditions (densities) *I*(0) = *I*_0_, *P*(0) = *P*_0_, *A*(0) = *A*_0_. We fit this model to three corresponding time series for each host to elucidate these interactions. We assume the virus replicates with a logistic growth rate *r*(1 − *P*(*τ*)/*K*), with a within-host virus carrying capacity absent adaptive immune response, *K*. The decision to model viral replication as logistic instead of exponential growth was made for two reasons. First, even in the hypothetical absence of an adaptive immune response, there is not an infinite number of cells to infect. Second, we use logistic growth to have a concentration-dependent viral replication rate, where the virus replicates more rapidly when fewer cells are infected. The innate response clears the pathogen at rate *θ*. The adaptive immune response, mediated by neutralizing antibodies produced by the host’s B cells, clears the pathogen at rate *δ*. Upon virus introduction, the adaptive immune response is activated via two pathways: It ramps up with a rate *b* independent of the innate response and responds to alarm signals induced by innate immune activation with a rate *aI*(*τ*)/(1 + *I*(*τ*)) (Tizard, 2017).

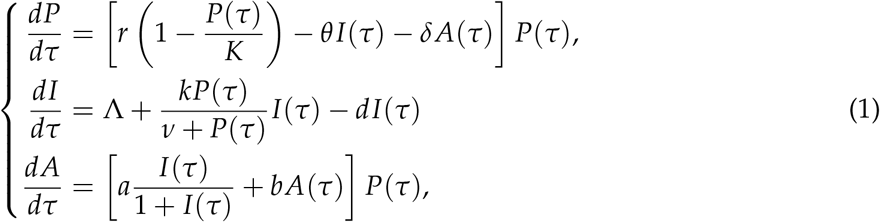

**Figure 1:**
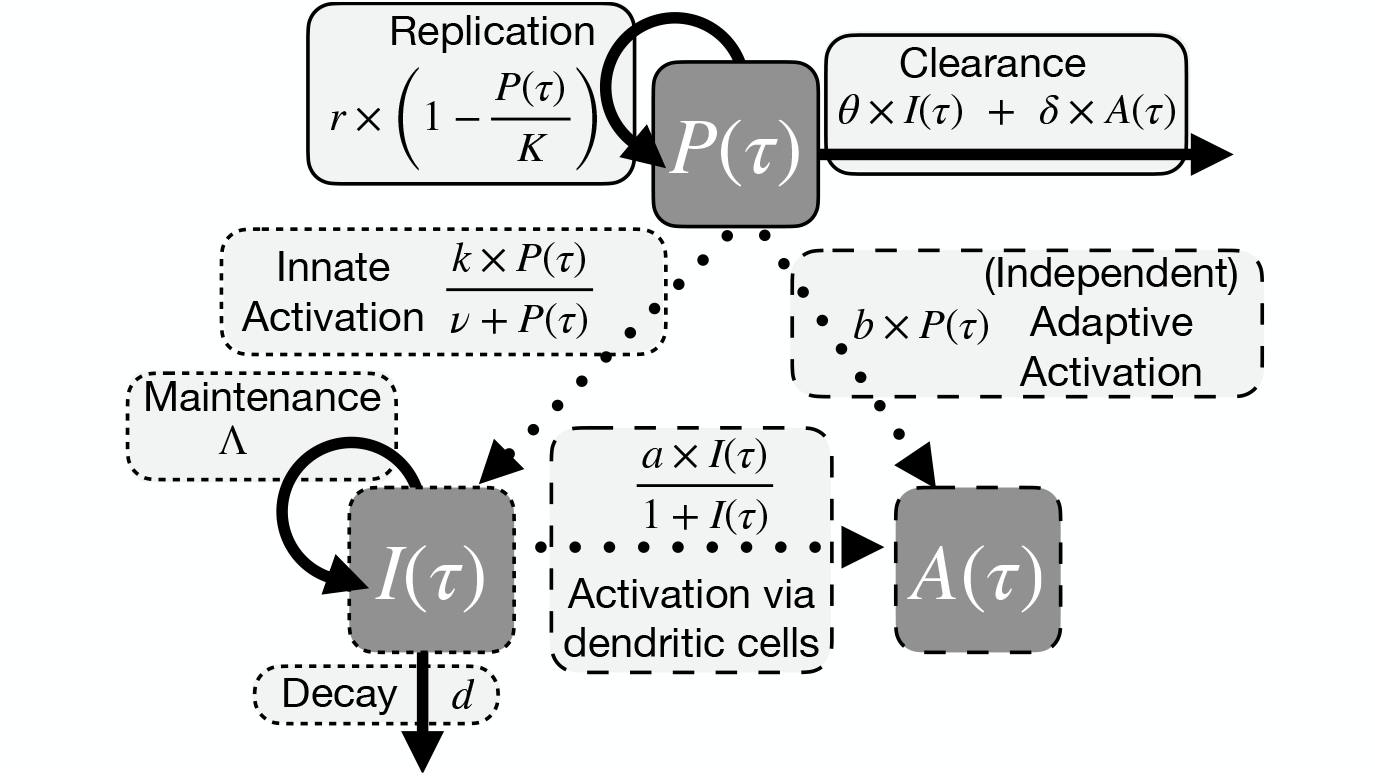
Model diagram corresponding to system (1), compartments are pathogen, *P*, innate immune response, *I*, adaptive immune response, *A*, and *τ* is time since infection began. Solid arrows represent population transitions (replication, clearance, maintenance, decay), and dotted arrows represent activation interactions. The border around arrow labels corresponds to the associated compartment: solid for pathogen, short dash for innate, and long dash for adaptive.

Innate immune responses, characterized by a marker of inflammation (haptoglobin), were assessed during experimental infection of buffalo with FMDVs, which yielded the most consistent fits compared to other measures of innate immunity that we assessed [see Supp. Material]. Our models assume that innate immune response effectors are always at a maintenance level Λ/*d* in the absence of a virus. Upon exposure to the virus, the inducible innate immune response is activated at maximum rate *k* with half saturation constant, *ν*, in terms of viral load, and decays at rate *d*, fixed at mean values inferred by Glidden et al. (2018).

Model parameters are identifiable given the data that we collected [Fig. 2, Supplementary figure S4]. In particular, we note that across 12 contact-infected hosts, the mean average relative error is well below the introduced noise level of 50%. This noise level was chosen so that our empirical data could have been plausibly generated by the mathematical model, barring a few obvious outliers (i.e., that the non-outliers fall within the 95% confidence intervals; see Figs. 2, S5, S6).

**Figure 2:**
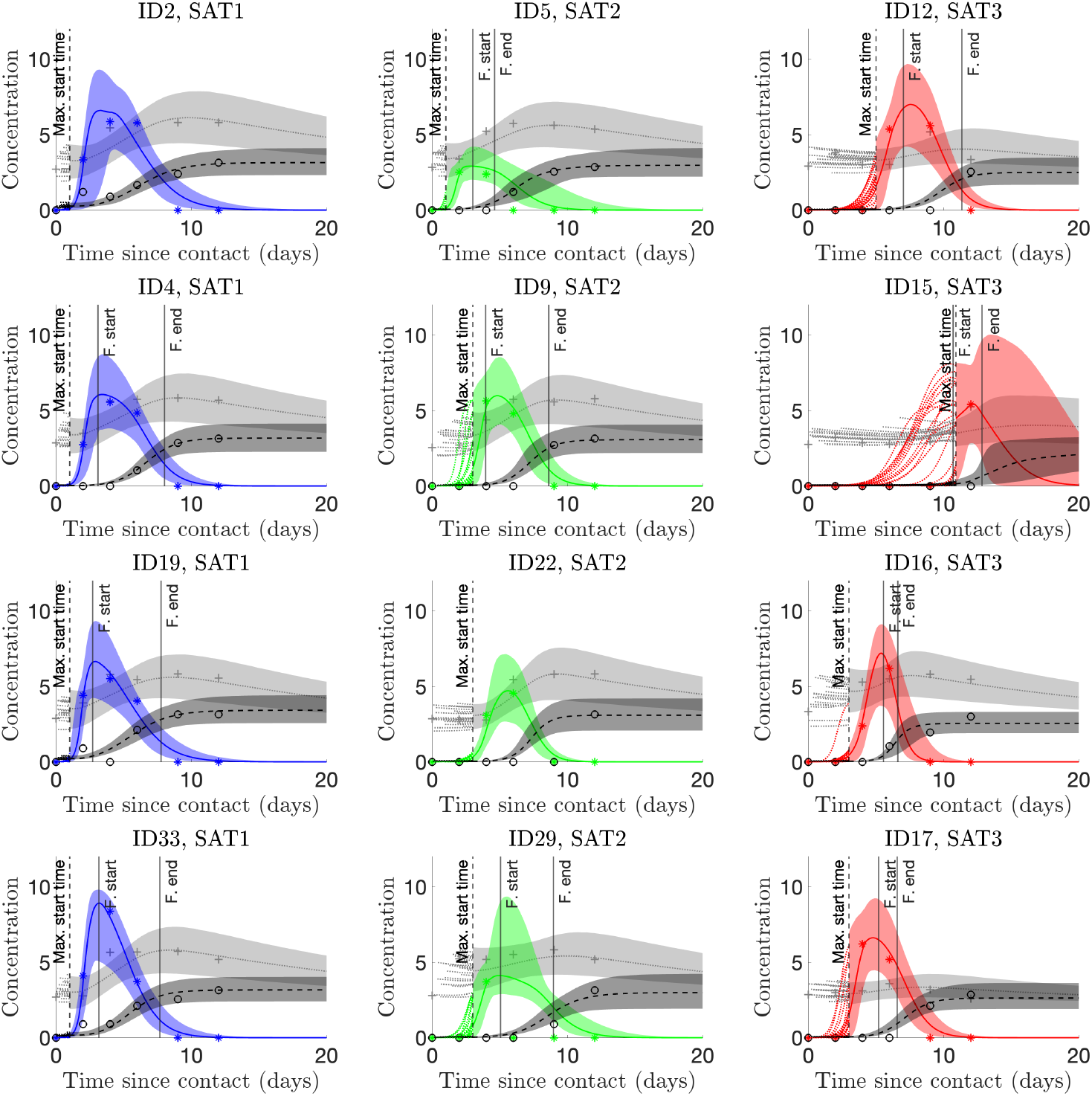
Within-host dynamics of FMDV in African buffalo. Our model reproduces *in vivo* observations. The vertical dashed line indicates the host-specific max infection start time (Gubbins, 2021). Before this line, the dotted lines are a sample of 20 model trajectories. Medians are represented by lines and 95% confidence intervals given by shades; Haptoglobin is measure of innate response (light grey) [log_10_(*μg*/*mL*)], viral data is represented by colored shades [log_10_(genome copies/mL)], and virus neutralization titer [log_10_(VNT)] is measure of adaptive response (dark grey). Virus data are indicated with asterisks, Haptoglobin with plusses, and VNT with circles. The ‘F. start’ and ‘F. end’ lines indicate the period a fever was detected. The lack of fever indicates logger malfunction. A value of zero for a given data point indicates that the data falls below the detection limit.

As an additional reality check for our models, we compared model output against data on fevers mounted by the animals. The model-inferred time course of immune and viral dynamics was reasonable in the context of clinical signs (see Perez-Martin et al., 2022, for details of the temperature data and the shaded boxes in Fig. 2 for a visual representation of these inferred fever quantities.) There was some variability among viral strains and individual hosts in the timing and magnitude of the fever response to FMDV infection. However, for the most part, buffalo mounted fevers as viral loads peaked and maintained elevated body temperature until most of the virus had been cleared – as one would expect for an acute viral infection (Tizard, 2017).

**Table 1:**
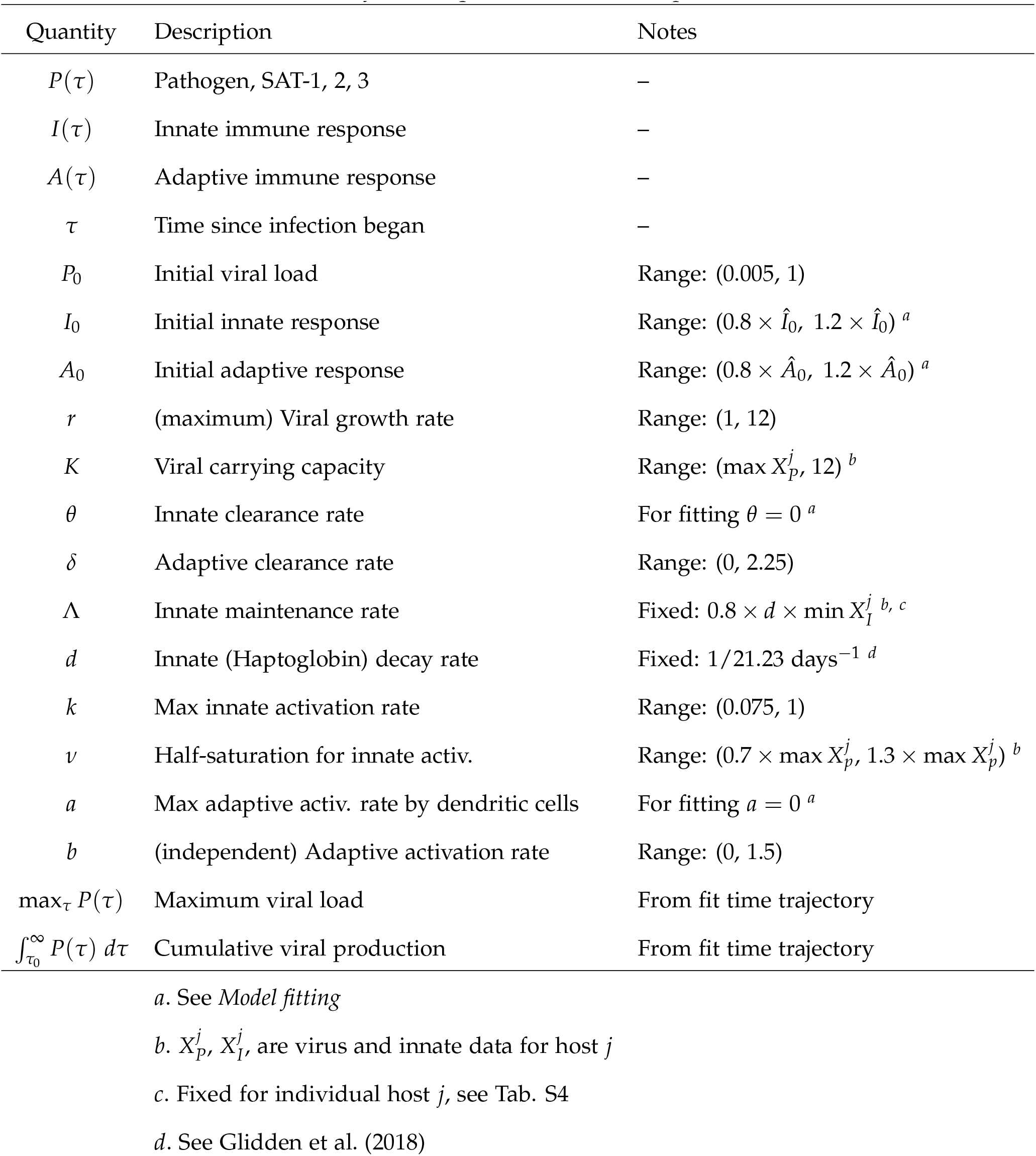
Key model quantities with description.

### Model fitting

Parameter estimates for each host were obtained by taking a weighted sum of the numerical solutions to the model compartments (with weights, which were treated as a chosen hyperparameter for each host so that average relative error was reduced while retaining the visual quality of model fits, see Macdonald et al., 2024) as an objective function in a simultaneous non-linear least squares fitting (via the interior reflexive Netwon-Rhapson method as implemented by MATLAB’s lsqcurvefit function) to three time-series each: innate response as measured by Haptoglobin [log_10_(*μg*/*mL*)], virus [log_10_(genome copies/mL)], and adaptive response as measured by virus neutralization titer [log_10_(VNT)]. This was done for each host given a wide range of possible host-specific infection start times as part of a profile (pseudo) likelihood approach. Inherent to this choice is that it is unclear how to pool the infection start times of each host for a hierarchical approach. While it is likely not strictly true that each host’s infection start times are independent, we cannot identify which host infected which host from a single infection experiment. Additionally, both serotype-specific transmission qualities *and* host-specific immunogenicity likely play a role in infection start time relative to the initiation of contact.

For both the viral and adaptive immune assays, an individual data value of 0 is an indication of failure of detection (due to concentration falling below the detection limit, see Perez-Martin et al., 2022). As such, when taking the log of the innate immune data, which was initially ecorded in *μ*g/mL, before fitting, we take the log of only the positive values to retain consistency with the other assays. Thus, in the model time trajectories, a value of zero indicates that the virus has been cleared from the blood after the virus’s proliferation period. We cannot know precisely when clearance occurs, but only when the virus is below detectable levels. Built-in errors such as this are a large part of why we perform practical identifiability analysis, as described below.

Data was collected for *contact-infected hosts* on contact days *t* = *{*0, 2, 4, 6, 9, 12, 28*}* and for *needle-infected hosts* on contact days *t* = *{*−2, 0, 2, 4, 6, 9, 12, 28*}*. For our data fitting, we first randomly draw an infection start time, *τ*_0_ ∈ (0, *τ*^*\**^ − 1) according to the posterior distributions obtained for each host in (Gubbins, 2021), where *τ*^*\**^ is the time of the first measured positive viral load in contact days. Once done we obtain initial estimates for *initial viral load, P*_0_, and *viral growth rate, r*, given *τ*_0_ and assuming a simple exponential growth model up to time *τ*^*\**^, that is for host *i*:

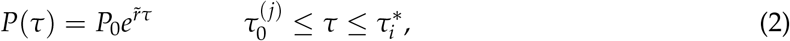

where 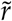 is the net viral growth rate (in the presence of innate immune response). There are two implicit biological assumptions inherent to this fitting. First, since the innate immune response is always present, estimating *r* this way will account for the killing of the pathogen by the innate response (thus providing a lower bound for *r*), and second before the presence of sufficient adaptive immune response pathogen growth will be exponential (instead of logistic). The second is a common assumption in the literature when estimating viral growth rates (Lloyd, 2001; Pawelek et al., 2012). This estimate of *P*_0_ was retained and fixed, while this estimate for viral growth rate was used as a lower bound and initial estimate for viral growth rate when simultaneously fit with other model parameters when we consider the following viral-immunological model (obtained by setting *θ, a* = 0):

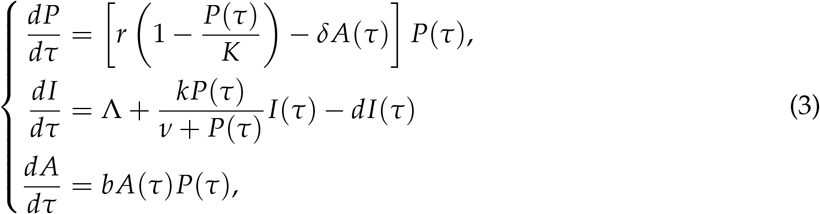

For fitting, we ultimately set *a, θ* = 0. This simplification is numerically justified in that when fit, these quantities were approximately *O*(10^−10^) (see program files, Macdonald et al., 2024). Setting *θ* = 0 does not indicate that adaptive clearance and innate clearance rates cannot be differentiated but that viral replication and innate clearance can not be separately identified. At some level, the innate immune response is always present *in vivo*. Thus, the estimate of model parameter *θ* ≈ 0 indicates the estimated viral replication rates being net viral replication rates in the presence of innate immune response.

For parameter *a* ≈ 0, a value of 0 does not indicate that this pathway is unimportant, but instead that we do not observe it in action. This lack of observation may be due to our data representing a single snapshot in time every two days or to the chosen proxies used to measure both adaptive and innate immune responses. We anticipate that if it were practicable to gather data from each host more frequently or if other proxies for the immune response were measured, then estimation of this parameter may be possible. However, collecting samples from each host requires sedation, and more frequent sampling is thus not practical.

To estimate *A*_0_, *I*_0_, we first fit a simple linear regression model to the first three time points of the adaptive and innate immune response levels, respectively, for each host. We then took the linear model prediction (denoted 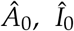) of the immune response at time *τ*_0_ as an initial guess for these quantities, and our model fitting allowed these parameters to vary between 80% and 120% of the regression prediction.

### Practical identifiability analysis and uncertainty quantification

To assess identifiability, confidence in our fitting procedure, and identify significant differences between (i) serotypes within a single route of infection and (ii) within a given serotype across routes of infection, we conducted uncertainty analysis via Monte Carlo simulations (10,000 individual replicates per host) simultaneously with our baseline profile likelihood estimation. This analysis was carried out in the following manner (see Miao et al., 2011; Tuncer and Le, 2018; Wieland et al., 2021, for detailed reviews of identifiability analysis for non-linear ODE models.)

On replicate *k*, for host *j* and drawn infection start time, *τ*_0,*j,k*_ we first obtain parameter estimates, 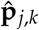 given the drawn infection start time and host. Next, we generate a new dataset using equation (4) under the assumption that the measurement error is independently and normally distributed with variation relative in magnitude to the expected total at each data point,*i*, 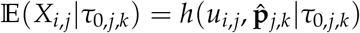:

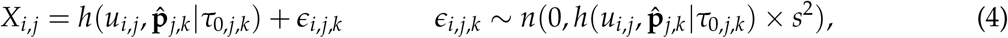

where *u*(*τ*) = [*I*(*τ*), *P*(*τ*), *A*(*τ*)]^*T*^.

Then, we refit the model (3) to the newly generated dataset according to the steps outlined in the preceding section. Finally, we calculate the relative error between model fit parameters from the baseline data and the generated data for each randomly drawn infection start time, host, and parameter. We then calculate the average relative error (ARE) for all model parameters across replicates for each host and the mean ARE across both needle and contact-infected hosts (see Fig. S4). Let *p*_*ℓ,j*_, *ℓ* = 1, …, *L* indicate an individual parameter for host *j*, and denote its ARE, sample mean ARE, and sample standard deviation as

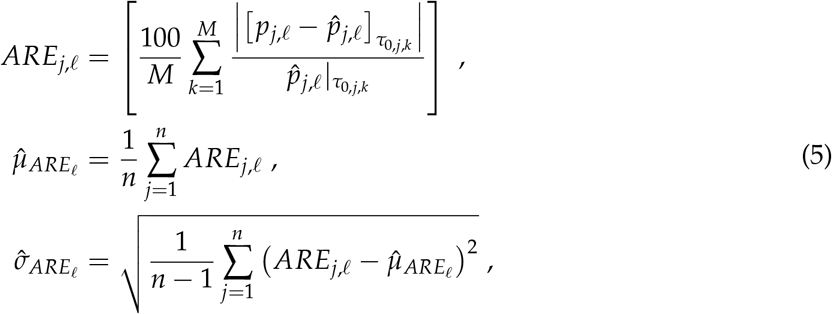

respectively. Following Tuncer and Le (2018); we say that parameter *p*_*ℓ*_ is (strongly) practically identifiable if the sum of its sample mean and standard error is less than the introduced noise level:

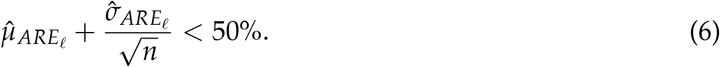

The results of this analysis indicate that for the contact infected hosts, the key model parameters and associated quantities are all practically identifiable assuming the 50% noise level in measurement error (that is *s* = 0.5 in equation (4); see figure 2 and Supplementary Figure S4).

## Results

### How do viral and immune dynamics vary among FMDV strains?

The three viral strains exhibited contrasting dynamics within buffalo hosts. Relative to the experiment start, SAT-1 attains its maximum viral load most rapidly, on average 3.14 days post-contact, while SAT-2 and SAT-3 took 4.438 and 7.22 days, respectively [SAT-1: 2.79-3.62 days; SAT-2: 3.95-4.89 days; SAT-3: 6.44-7.95 days; Fig. 3c]. Indeed, the viral growth rate was negatively correlated to time to maximum viral load, latency period, and infection start time (in contact days) among individual hosts [Fig. 3d, i and Fig. 5a Pearson correlation across median parameter estimates from 10,000 Monte Carlo simulations for each of the 12 sample hosts *ϱ* = −0.89, *p* = 8.5 *×* 10^−^5; *ϱ* = −0.83, *p* = 8.2 *×* 10^−4^; and *ϱ* = −0.85, *p* = 5.0 *×* 10^−4^ respectively]. Finally, viral replication and initial viral load are tightly positively correlated *ϱ* = 0.97, *p* = 2.1 *×* 10^−7^. These relationships point to life history variation among viral strains. Viruses with fast growth rates appear to reach their within-host peak load quickly after contact initiation but require comparatively more viral material to mount a successful infection.

**Figure 3:**
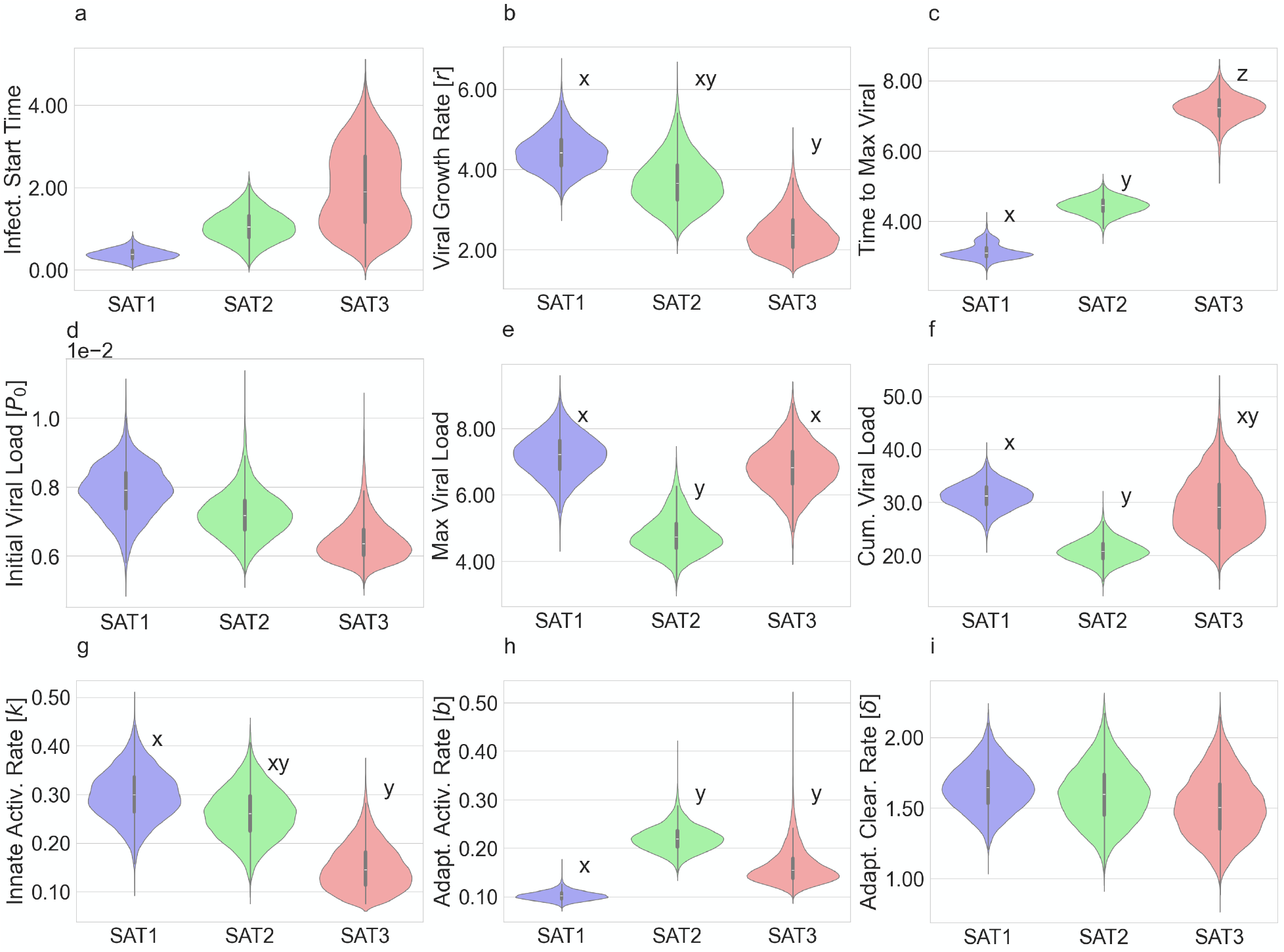
Variation among FMDV serotypes in viral and immune parameters defining within-host dynamics. Violin plots represent the empirical distribution of serotype sample mean parameter estimates generated from bootstrap and identifiability analysis procedure; compact letter display is used to denote statistically significant differences as indicated by the 95% CI for the difference between means not containing zero.

Host immune responses to FMDV infection also varied by strain. Buffalo activated innate immune responses [*k*] (as measured by haptoglobin, an acute inflammatory protein) more rapidly when infected with SAT-1 than SAT-3, while SAT-2 is intermediate between them [SAT-1: 0.30, *95% CI* 0.200-0.404; SAT-2: 0.261, *95% CI* 0.162-0.365; SAT-3: .150, *95% CI* 0.0773-0.256, Fig. 3g]. Buffalo activated adaptive immune responses, as measured by FMDV neutralizing antibody titer [*b*], more rapidly for SAT-2 than either SAT-1 or SAT-3 [SAT-1: 0.102, *95% CI* 0.085-0.123; SAT-2: 0.221, 0.174-0.279; SAT-3: 0.164, 0.116-0.263]. Finally, SAT-1 hosts had significantly (*α* = 0.05) elevated adaptive response levels at the start of infection relative to SATs 2,3 [SAT-1 .187, 0.132-0.261; SAT-2 0.048, 0.034-0.063; SAT-3 0.047, 0.033-0.062] Collectively these results are indicative of variation among hosts in the speed of their immune responses [Figs. 3,4].

**Figure 4:**
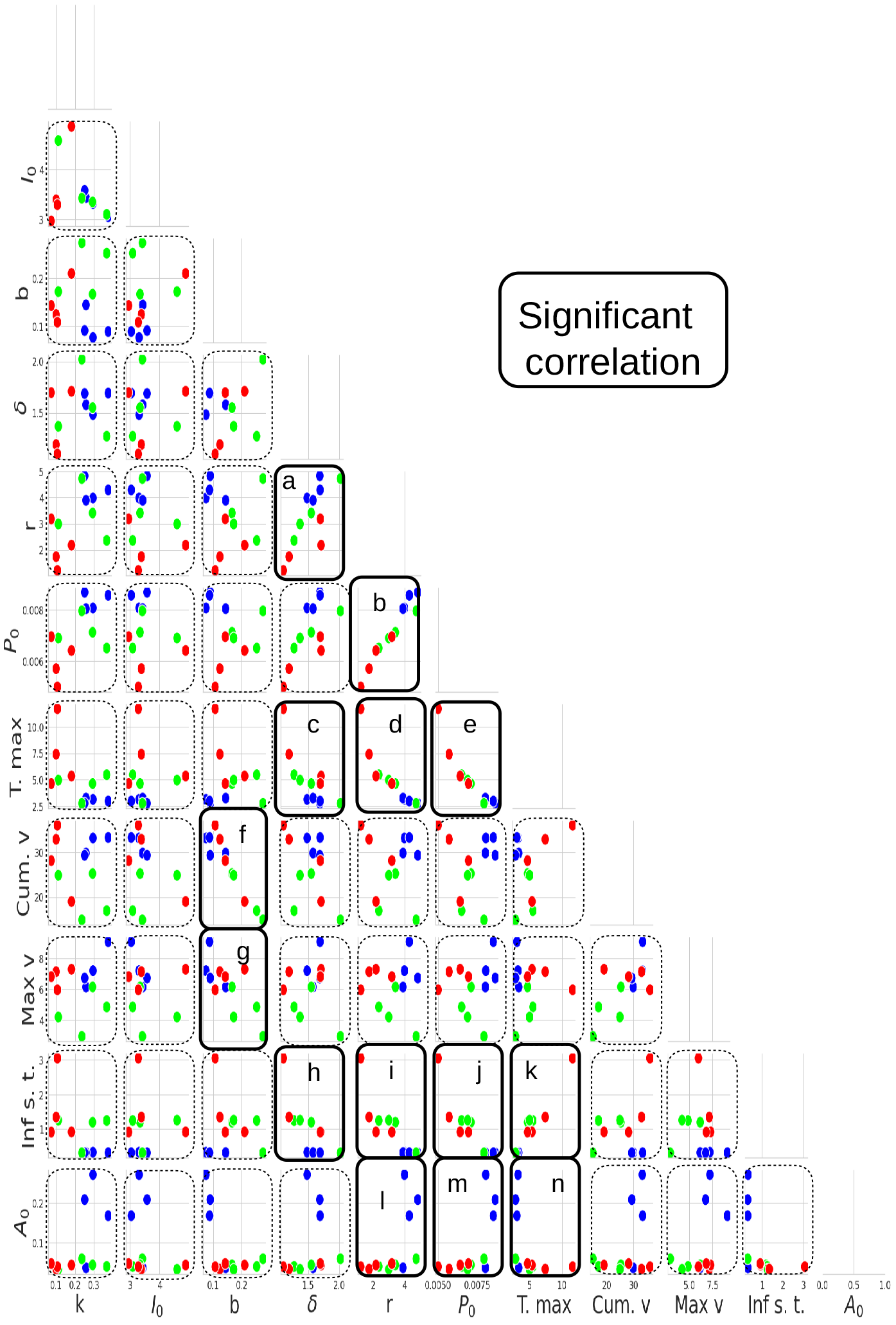
Co-variation between parameters capturing viral and immune dynamics within buffalo hosts. Blue indicates SAT-1, green SAT-2, red SAT-3. Scatter plots show median parameter estimates from 10,000 replications for each host generated by Monte Carlo simulations. Pairs of parameters that have significant Pearson correlations are framed in solid black lines.

### Which viral & immune parameters determine viral fitness within hosts?

The observed differences in viral dynamics and host responses to infection resulted in differences in viral fitness among strains: SAT-1 attained high maximum and cumulative viral titers in buffalo hosts. SAT-2 lagged conspicuously behind the other strains in cumulative and maximum viral load. SAT-2’s maximum viral load was 67.3% [95% CI 50.3-90.9%] that of SAT-1 and 71.1% [95% CI 51.8-98.3%] that of SAT-3’s; and its cumulative viral load was 32.77% [95% CI 13.9-48.25 %] lower than the fittest strains (SAT-1, SAT-3) [Fig. 3e,f]. Total viral production by each host (cumulative viral load) was driven overwhelmingly by variation among buffalo in adaptive activation rate: a more rapid adaptive immune activation rate was associated with lower cumulative viral load [Fig. 4f; *ϱ* = −0.95, p = 3.6 *×* 10^−6^]. Within-host viral success thus appeared constrained by each strain’s ability to evade host immune responses rather than the viruses’ replication capacity. These findings suggest that different viral life histories can result in similar fitness in cumulative and maximum viral loads mediated by the viral interaction with the host’s immune responses.

### How do within-host dynamics relate to virus transmission among hosts?

To explore how within-host FMDV dynamics might scale up to affect viral transmission among hosts, we compared parameters fit to our within-host model to population-scale parameters derived from the same set of experiments (Jolles et al., 2021). These analyses indicate that variation in viral growth rate among buffalo correlates tightly and negatively with latent period [Fig. 5a], such that buffalo harboring fast-growing viral populations progress to the infectious class more rapidly. Further, viral growth rate within the host may correlate with variation among viral strains in transmission rates [Fig. 5b]: Fast-growing SAT-1 transmitted among buffalo most readily, followed by SAT-2, while SAT-3’s slower-paced time to within-host maximum was matched by slower transmission among hosts. Variation among hosts in the rate at which adaptive immune responses against FMDV were activated correlated with each host’s infectious period [Fig. 5c].

**Figure 5:**
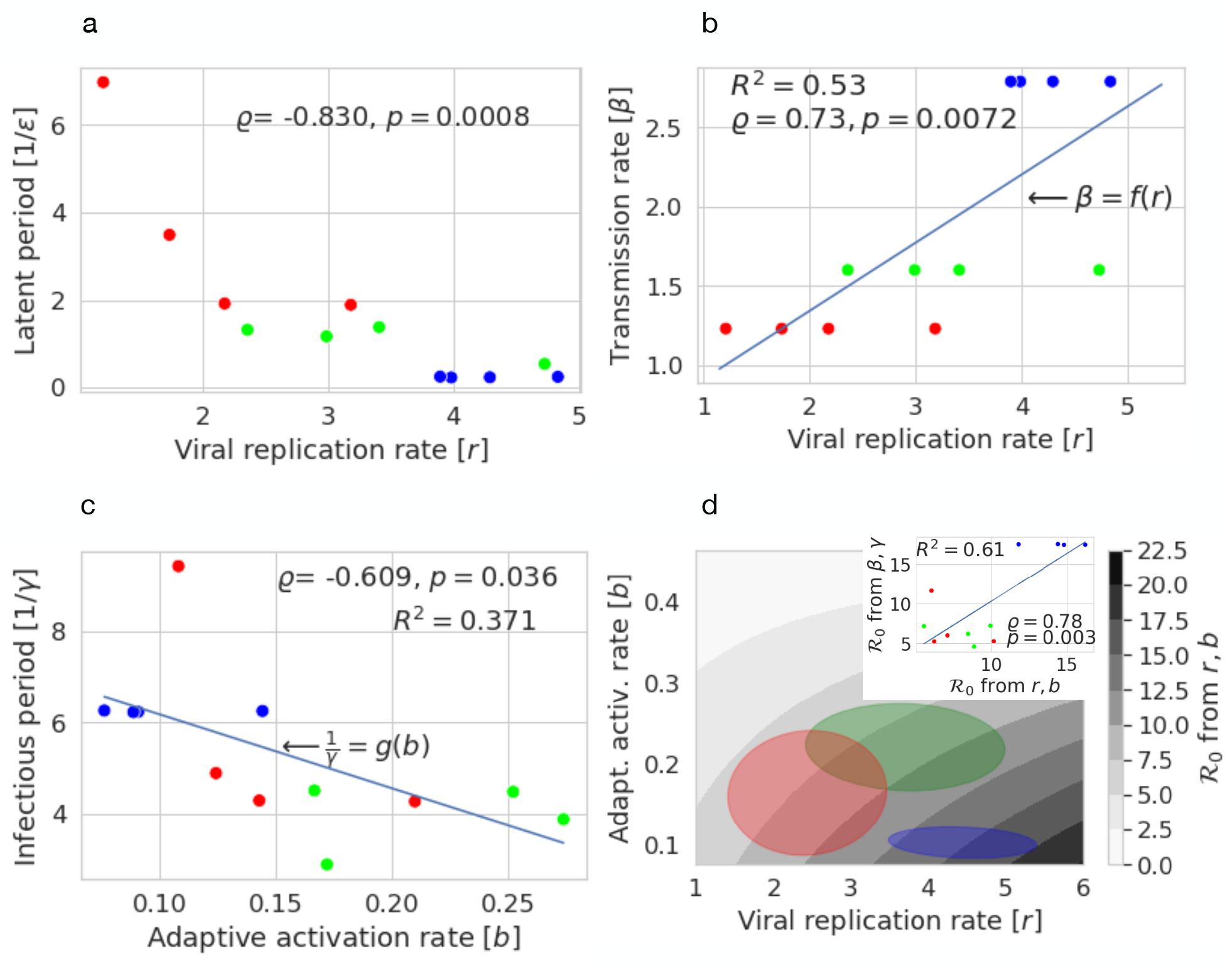
Consilience across scales: (a) Viral growth rate is negatively correlated with the latent period (b) sample mean viral growth rate has apparent positive trend with transmission rate, the blue line is the best fit linear function, *f* (*r*) (c) adaptive activation rate is negatively correlated with the infectious period, the blue line is the best fit linear function, *g*(*b*) (d) ℛ_0_, a population level quantity which summarizes transmisibility, may be viewed as a function of viral growth rate and adaptive activation rate. The contour is generated by taking *f* (*r*) *× g*(*b*). (inset) Correlation between within-host and between host parameter estimates for ℛ_0_. In the between host estimates, the group estimate of transmission rate is used; all other parameters are individual host estimates. Blue indicates SAT-1, green SAT-2, red SAT-3.

Together, the transmission rate and infectious period determine the relative basic reproductive number (ℛ_0_) of the virus, which can be calculated as ℛ_0_ = (transmission rate)*·*(infectious period) given constant contact rates and host density (reasonable assumptions given our experimental setup). Our data on cross-scale linkages in viral dynamics suggest that viral invasion potential may be predictable from within-host dynamics. We explored this idea by comparing ℛ_0_ established previously (Jolles et al., 2021) by observing transmission of FMDV between hosts, with ℛ_0_ calculated based on within-host viral and immune kinetic parameters: We estimated transmission rate and infectious period as increasing and decreasing linear functions of viral growth rate and adaptive activation rate, respectively, and computed ℛ_0_ as their product [see Fig.5b, c].

The immune parameters used in this estimate (*r,b*) are from this work, and the population scale parameters (1/*γ*, 1/*ϵ, β*) are from (Jolles et al., 2021). The data points in Fig. 5 represent median parameter estimates for each contact-infected host. The infectious and latent periods were both assumed to be gamma distributed, with posterior means for the latent period: 0.5 days [95% CI 0.02-2.4], 1.3 days [0.1-3.5], 2.8 days [0.5, 7.0] and posterior shape parameters 1.2 [0.1-8.7], 1.6 [0.2-9.2], 1.6 [0.2-8.3] for SAT-1, 2 and 3 respectively (Jolles et al., 2021). Similarly, for the infectious period, posterior means were estimated at 5.7 days [4.4-7.4], 4.6 days [3.5-6.3], and 4.2 days [3.2, 5.8], and posterior shape parameters 11.8 [3.5-33.5], 8.7 [2.4-27.0], 11.8 [3.3-35.3].

We then compared these basic reproductive numbers derived from within-host viral dynamic parameters [Fig. 5d] to reproductive numbers estimated by observing transmission among hosts in our experiments (Jolles et al., 2021), and found that both estimates of ℛ_0_ match qualitatively [Fig. 5d inset]. Our data on three strains of FMDV demonstrate a good match between ℛ_0_ estimated from within-host parameters with ℛ_0_ measured by observing disease transmission among hosts. However, more than three strains of FMDV will need to be studied to test the robustness of this finding.

## Discussion

In this study, we used time series data from viral challenge experiments involving 12 buffalo acutely infected with three strains of FMDVs (4 buffalo per strain) to parameterize a mathematical model capturing the dynamics of viral growth and its curtailment by the host’s immune responses. We sought to determine: (i) How do viral and immune dynamics vary among FMDV strains?; (ii) Which viral & immune parameters determine viral fitness within hosts?; and (iii) How do within-host dynamics relate to virus transmission among hosts? Despite the moderate number of host individuals included in the study, identifiability analysis of our model parameters suggests that our models capture the data well: Identifiability analysis yielded robust interval estimates for model parameters by replicating the sample 10,000 times as well as a metric (average relative error) to assess whether or not model parameters can be reliably identified from available data. Here, we found that our model parameters were practically identifiable at an introduced noise level of 50% (chosen to cover the original data, see Figs. S4-S6) and that our results correspond well with independently estimated metrics of fever within the same hosts (see Fig. S4).

Our results uncover variation among viral strains in within-host dynamics. SAT-1 was able to transmit most rapidly, leading to an early peak in viral load just 3-4 days after the start of contact. SAT-2 achieved its maximum viral load somewhat more slowly, reaching its peak about a day later than SAT-1. By contrast, SAT-3 appeared to follow a strikingly different strategy, attaining its peak viral load three days later than SAT-1. However, interestingly, despite the significant disparity between SAT-1 and SAT-3 in the time course of viral proliferation within the host, within-host fitness of these strains was quite similar: SAT-1 and SAT-3 attained similar maximum and cumulative viral loads. Both measures showed that SAT-2 had sharply reduced fitness compared to the other two strains - despite kinetic similarities with SAT-1. This contrast appeared related to differences in host immune responses to the three strains. SAT-2 elicited more rapid and effective innate and adaptive immune responses than SAT-1 and SAT-3. Differences in viral production among individual hosts were thus mediated by variation in viral interactions with the host’s immune responses: fast, effective adaptive immune responses limited cumulative viral production within a given host. Rapid activation of adaptive immune responses also curtailed each host’s infectious period; thus, the host’s antibody responses shut down viral production and the potential for transmission to other hosts. These findings suggest that different viral life histories – characterized by variation in latency period and time to maximum population size within the host relative to contact days – can result in similar viral fitness in terms of the amount of virus produced in an individual host. Indeed, adaptive immune activation rate and cumulative viral production were so tightly correlated [*ϱ* = −0.93] as to be practically synonymous, indicating that viral fitness as measured by cumulative production in an individual host near exclusively reflected how speedily the host was able to mount a neutralizing antibody response to infection. At least within the parameter space defined by the viral strains and buffalo that we worked with, no other parameters of the virus-host interaction played a significant role in determining the viral production of each host.

On the other hand, acute transmission rates, the per day expected number of successfully infected contacts of the three strains, estimated in previous work (Jolles et al., 2021), appear to follow the variation in median viral replication rate per serotype and not viral load. SAT-1 had the most rapid time to peak load and transmission rate, SAT-2’s were intermediate, and SAT-3’s were the lowest, whereas viral load patterns did not match the transmission rate variation among strains. These observations are based on a sample size of three strains: unlike our other trait associations, which evaluated variation in viral and immune dynamics across 12 hosts, our estimates of viral transmission rate are group averages. This difference arises because we cannot distinguish which individual hosts transmitted infection during our experiments - we merely recorded the timing of new infections in each group. Whole genome sequencing of the virus recovered from each buffalo during the experiments might allow us to pinpoint who infected whom, elevating the precision of our transmission rate estimates to the individual level. However, even for a rapidly evolving RNA virus such as FMDV, genomic differentiation of experimental strains during a single transmission cycle from needle-infected to in-contact hosts may not prove sufficient to identify donor and recipient hosts confidently. Finally, a greater number of viral strains would ideally need to be studied to assess the generality of our findings.

Two additional observations are consistent with the finding that viral transmissibility appears to be related to the within-host replication rate. We found that infection start time, latency period, and time to maximum viral load negatively correlated with viral growth rate. In our results, high transmissibility is correlated with high viral growth rates and the ability to (apparently) produce large quantities of virus. In essence, a high viral growth rate in the blood may be reflected in high production in mucosal cells and, therefore, shedding of virus. Variation in pathogen contagiousness is often related to differences in the infectious dose sufficient to cause infection in a new host (Fine, 2003) – indeed, FMDVs can notoriously transmit with tiny amounts of inoculum. Just a few virions suffice to propagate infections of some FMDVs to susceptible hosts (Alexandersen et al., 2003; Quan et al., 2004), contributing to these pathogens’ hallmark contagiousness. Similarly, short incubation periods can contribute to the rapid spread of highly transmissible pathogens (Fine, 2003; Grassly and Fraser, 2008; Nishiura et al., 2020; Sartwell, 1995). As such, these observations lend credence to the idea that aggressive within-host replication in FMDVs may indicate high transmission capacity among hosts.

It is important to note that our findings linking viral growth and transmission rates refer specifically to transmission during acute infection. In addition, FMDVs can be transmitted from carrier buffalo that retain the virus in follicular dendritic cells of the palatine tonsils (Juleff et al., 2008) long after the virus has been cleared from the blood. For the FMDV strains we studied, we previously estimated that SAT-1 and SAT-3 transmit from carrier hosts at much reduced (approx. two orders of magnitude lower) rates compared to transmission from acutely infected hosts, and carrier transmission of SAT-2 is even rarer if it occurs at all (Jolles et al., 2021).

We note that serotypes are defined by how they interact with the host immune system Tizard (2017). Thus, differing selection pressures on the three strains may explain these differences in the persistence mechanism for SAT-1 and SAT-3 vs SAT-2. Though SAT-1, 2, and 3 are not spatially segregated, their occurrence among host species may vary (Jolles et al., 2021; Sirdar et al., 2021). Thus, while local adaptation due to spatial segregation is unlikely, there may be serotype-specific adaptation to different dominant host species. For example, SAT-2 is the most common strain causing outbreaks in cattle (Blignaut et al., 2020). Nonetheless, at least for SAT-1 and SAT-3, viral transmission from carrier hosts may play a crucial role in maintaining the long-term persistence of these viruses in buffalo populations by sparking epidemics in newly susceptible calf cohorts. In contrast, the mechanisms that sustain SAT-2 endemic persistence at the herd and landscape scales are uncertain, with carrier transmission alone appearing insufficient to maintain it between birth pulses. For this strain, mechanisms such as antigenic shift, loss of immunity, or spillover among host populations may be necessary to explain persistence (Jolles et al., 2021). Future work could use phylodynamic approaches (Volz et al., 2013) to evaluate the viral transmission and divergence among wildlife and domestic animal hosts.

Bringing together two key findings – that adaptive immune activation drives the duration of the infectious period in each host, and viral growth rate may determine the acute viral transmission rate among hosts – we estimated ℛ_0_, the virus’ relative basic reproductive number from within-host dynamic parameters. We showed that estimates based on viral replication rate and adaptive activation rate qualitatively matched ℛ_0_ estimates previously derived from observed transmission events among experimentally infected and naive buffalo (Jolles et al., 2021), suggesting that viral invasion potential may be predictable from within-host dynamics. With just three viral strains and 12 host individuals to work with, these findings are necessarily tentative. We assumed linear relationships linking viral replication and transmission rates and adaptive activation rate with the infectious period, when in fact, these functions might follow more complex shapes, and our power to evaluate how well ℛ_0_ estimated from within vs. among host processes match, is limited. Nonetheless, the possibility of predicting pathogen behavior in host populations from within-host experiments is tantalizing: studying pathogen strains in individual animals is far more tractable than investigating their behavior at the population scale, yet predicting which pathogen strains are likely to spread and persist in host populations of interest is an urgent priority in the face of globally accelerating pathogen emergence (Cleaveland et al., 2007).

Our study illustrates the value of taking a functional approach to understanding the consequences of viral diversity. By documenting variation among viral strains in terms of their life history traits, rather than focusing on genomic variation, we were able to bridge biological scales from kinetics within individual animals to transmission among hosts. This was effectively an information reduction step - zooming out to extract relevant life history signals to understand viral dynamics across scales. Future work could explore whether a functional approach to viral dynamics can be extrapolated down, leveraging in vitro studies of viral kinetics to predict viral life history traits and interactions with the host, and should test the generality of our findings by expanding the number of viral strains that are included. This would also allow parameterization of multi-scale models that explicitly link within-host and between-host dynamics and model validation through evaluating viral dynamics and population structure in natural host populations.

## Acknowledgments

We thank the South African Parks Veterinary Wildlife Services and State Veterinary Services staff at Skukuza for their help with animal capture and sample collection. Lab work and fieldwork were completed by H. Combrink, C. Coon, C. Couch, E. Devereux, B. Dugovich, K. Forssman, C Glidden, J. Masseloux, A. Sage, D. Sisson, H. Tavalire, and D. Trovillion. Ethical clearance was obtained from Oregon State University (ACUP 4478), South African National Parks (project no. OLAE 1157), the South African Department of Agriculture, Forestry and Fisheries: Directorate of Animal Health (Section 20 permit 12/11/1/8/3), and Onderstepoort Veterinary Research Animal Ethics Committee (100261Y5). JCM and HG are supported by a US NSF RAPID grant (no. DMS-2028728) and NSF grant (no. DMS-1951759), JCM by a Zuckerman Postdoctoral Scholarship, and HG by a grant from the Simons Foundation/SFARI 638193. Experimental work and epidemic scale model development performed by AJ, BB, EG, E.P-M, and SG was supported by USDA-NIFA AFRI grant 2013-67015-21291 and by the UK Biotechnology and Biological Sciences Research Council grant BB/L011085/1 as part of the joint USDA-NSF-NIH-BBSRC Ecology and Evolution of Infectious Diseases program. SG was supported by UK Biotechnology and Biological Sciences Research Council grants BBS/E/I/00007030, BBS/E/I/00007036, and BBS/E/I/00007037. Ongoing work by all authors is supported by the joint NSF-NIH-NIFA Ecology and Evolution of Infectious Disease program (grant number 2208087).

## Statement of Authorship

JCM Software, formal analysis, mathematical modeling methodology, data & software curation, wrote the original draft. HG Conceptualization, mathematical modeling methodology, formal analysis, data & software curation, supervision. BB Experimental design, experimental data collection, and formal analysis EG Experimental data collection and formal analysis SG Formal analysis, software, mathematical modeling methodology EPM Experimental data collection and formal analysis AJ Conceptualization, experimental design, data collection, formal analysis, and supervision. All authors reviewed and edited the manuscript prior to submission.

## Data and Code Availability

All software and data for this project is available on [GitHub](https://github.com/MacdonaldJoshuaCaleb/FMDV-Fitting-Bootstrap) and is stored in cite-able format in the Zenodo repository (Macdonald et al., 2024). Software to generate posterior chains used to draw infection start times is available from Gubbins (2021). Population scale data was published along with our prior work (Jolles et al., 2021). Detailed computational methods are provided in the SI of this work. All materials are provided under a CC-BY-NC (non-commercial) 4.0 license.

## Notes

https://doi.org/10.5281/zenodo.10720079

### Competing Interest Statement

The authors have declared no competing interest.

### Summary of Updates

We have updated our approach to fitting initial conditions for each model compartment. We have also updated all figures, added new text including expanded discussion to the primary text. We have also added a new table and two new figures to the SI. Finally, we have added version information to the associated code repository and edited all software files for clarity.

