## Supplementary Materials for "Within-host viral growth and immune response rates predict FMDV transmission dynamics for African Buffalo"

**Joshua C. Macdonald** [1,2]

Hayriye Gulbudak[1,*]

Brianna Beechler[3]

Erin E. Gorsich[4]

Simon Gubbins[5]

Eva Pérez-Martin[5]

Anna E. Jolles[3,6,*]

1. Department of Mathematics, University of Louisiana at Lafayette, Lafayette, Louisiana, USA;

2. Current address, School of Zoology, Faculty of Life Sciences, Tel Aviv University, Tel Aviv-Yafo, Israel;

3. Carlson College of Veterinary Medicine, Oregon State University, Corvallis, Oregon, USA;

4. The Zeeman Institute for Systems Biology & Infectious Disease Epidemiology Research and School of Life Sciences, The University of Warwick, Coventry, UK

5. The Pirbright Institute, Woking, Surrey, UK

6. Department of Integrative Biology, Oregon State University, Corvallis, Oregon, USA

Supplementary Information for Viral growth and immune response rates predict FMDV transmission dynamics

### Contents

**S1 Model dynamics** 1

**S2 Discussion of model fitting approach** 1

**S3 Detailed description of Non-linear least-squares fitting** 2

**S4 Practical identifiability analysis and interval estimates** 3

**S5 Effect of route of infection on within-host viral and immune dynamics** 3

**S6 Supplementary Tables & Figures** 4

S3 Our model reproduces empirically observed *needle infected* viral and immune dynamics well. Median time trajectories are represented by lines and 95% confidence intervals by shaded regions; blue indicates SAT1, green SAT2, and red SAT3. Haptoglobin is used to measure the innate response (light grey shade) [$\log_{10}(\mu g/mL)$], FMDV viral data is represented by colored shades [$\log_{10}$(genome copies/mL)], and virus neutralization titer [$\log_{10}$(VNT)] is used as a measure of adaptive response (dark grey shade). Virus data are indicated with asterisks, Haptoglobin with plusses, and VNT with circles. . . . . . . . 8

S4 Identifiability analysis, (a) mean ARE for contact infected hosts with standard error, all model parameters are practically identifiable; (b) mean ARE for needle infected hosts with standard error, all parameters except initial viral load [$P_0$] are practically identifiable . . . . . . . . . . . . . . . . . . . 9

S5 The choice of 50% noise covers the the empirical contact infection data barring clear outliers (that is $s = 0.5$ in equation (4) of the main text). The points connected with lines are the empirical data. The dotted lines are a simple random sample of 500 time solutions corresponding to fits to data generated by equation (4) in the main text. Virus data are indicated with asterisks, Haptoglobin with plusses, and VNT with circles. . . . . . . . . . . . . . . . . . . . . . . . . . . . . . . . . . . . . . . . . . . 10

S6 The choice of 50% noise covers the the empirical needle infection data barring clear outliers (that is $s = 0.5$ in equation (4) of the main text). The points connected with lines are the empirical data. The dotted lines are a simple random sample of 500 time solutions corresponding to fits to data generated by equation (4) in the main text. Virus data are indicated with asterisks, Haptoglobin with plusses, and VNT with circles. . . . . . . . . . . . . . . . . . . . . . . . . . . . . . . . . . . . . . . . . . . 11

S7 Pearson correlation between sample median values for 12 contact infected hosts . . . . . . . . . . . . 12

#### S1 Model dynamics

Here we show that the adaptive immune response saturates in time.

$$\begin{cases} \dfrac{dI}{d\tau} = \Lambda + \dfrac{kP(\tau)}{\nu + P(\tau)} I(\tau) - dI(\tau) \\ \dfrac{dP}{d\tau} = \left[ r \left( 1 - \dfrac{P(\tau)}{K} \right) - \theta I(\tau) - \delta A(\tau) \right] P(\tau), \\ \dfrac{dA}{d\tau} = \left[ a \dfrac{I(\tau)}{1 + I(\tau)} + bA(\tau) \right] P(\tau), \end{cases} \tag{1}$$

*Theorem.* If $I_0 > 0$ or $A_0 > 0$, then the pathogen (within-the host) eventually clears ($\lim_{\tau\to\infty} P(\tau) = 0$), and subsequently the innate immune response antibodies decay and the adaptive immune response antibodies increase to a steady-state, respectively; i.e. $\lim_{\tau\to\infty} I(\tau) = \Lambda/d$ and $\lim_{\tau\to\infty} A(\tau) = A^+$, where $A^+ > 0$ depends on the initial condition; i.e. $A^+ = z(P_0, I_0, A_0)$.

*Proof.* Let $P_0 > 0$. By the second equation in (1), we obtain

$$P(\tau) \le P_0 e^{\int_0^\tau [r - \delta A(\tau)] ds}. \tag{2}$$

Without loss of generality, assume that $I_0 > 0$. Then $\exists \tilde{\tau} : A'(\tau) \ge 0$ for all $\tau \ge \tilde{\tau}$. Therefore $A(\tau)$ increases for all $\tau \ge \tilde{\tau}$.

Case (i) Assume $A(\tilde{\tau}) > \frac{r}{\delta}$. Then, we obtain $A(\tau) > \frac{r}{\delta}$, for all $\tau > \tilde{\tau}$. Therefore as $\tau \to \infty$, the RHS of the inequality (2) goes to zero. Then by comparison principle, we obtain $\lim_{\tau\to\infty} P(\tau) = 0$. Thus $\lim_{\tau\to\infty} I(\tau) = \Lambda/d$. Then $A(\tau)$ saturates as $\tau \to \infty$; i.e $\lim_{\tau\to\infty} A(\tau) = \bar{A}$, for some $\bar{A} > 0$, depending on the initial condition $(P_0, I_0, A_0)$.

Case (ii)Now suppose that $A(\tilde{\tau}) < \frac{r}{\delta}$. Assume that there exists $\hat{\tau}$: $P(\hat{\tau}) = 0$. Then we obtain that $P(\tau) = 0$, for all $\tau > \hat{\tau}$ and $\lim_{\tau\to\infty} I(\tau) = \Lambda/d$ and $\lim_{\tau\to\infty} A(\tau) = \bar{A}$, for some $\bar{A} > 0$. Now assume that $P(\tau) > 0$, for all $\tau > 0$. Then $A'(\tau) > 0$, for all $\tau > 0$. Hence there exists $\tau^+ > 0 :\ A(\tau^+) > \frac{r}{\delta}$. The rest of the proof follows the argument in case (i), completing the proof. □

#### S2 Discussion of model fitting approach

The model was fit for each host. For each host and over 10,000 replications for that host, an infection start time was drawn from the host-specific empirical distributions generated in our previous work (Jolles et al., 2021). We next obtained a point estimate for the model parameters given this drawn infection start time, as discussed in the primary text. This approach allowed the calculation of average relative error for each host and each parameter for that host to assess practical identifiability and generate interval estimates for the serotype sample mean parameters across a wide range of (combinations of) individual host infection start time. This amounts to a profile (pseudo) likelihood approach.

We have taken this approach, which is not hierarchical (Bayesian), for several reasons. First, the approach taken is more compatible with practical identifiability analysis and is a standard approach used when conducting such analyses (see Miao et al., 2011; Tuncer and Le, 2018; Wieland et al., 2021). Practical identifiability depends not only on the type of data collected but also on built-in errors such as detection thresholds and frequency of data collection. By fitting each host, we were thus able to assess more comprehensively the effect of individual variation among hosts on the identifiability of the model parameters. The relatively small sample sizes in this work (4 hosts per serotype) means that the distribution of the sample mean parameter values is not necessarily approximately normally distributed; there is a relatively large number of parameters that needed to be simultaneously estimated (twelve), and due to the nature of the contact infected experiments a precise infection start time is both unknown and host specific. These issues complicate the computation of accurate interval estimates, and in such circumstances, profile likelihood approaches are common (Wieland et al., 2021).

It may be reasonable to assume that virus-specific parameters such as replication rate are differentiated by serotype (and thus pooled) and most strongly determined by variation in life history characteristics intrinsic to the virus. However, this is not necessarily true for host-specific parameters such as the adaptive and innate immune response activation and clearance rates. Indeed, we considered it possible that, at least for some parameters, individual variation would provide a better explanation for differences among hosts than serotype-specific effects. Thus, we did not desire to make a priori assumptions about the relative importance of individual variation versus serotype-specific variation.

#### S3 Detailed description of Non-linear least-squares fitting

Consider the model

$$\begin{cases} \frac{d\mathbf{u}}{d\tau} = f(\tau, \mathbf{u}(\tau, \mathbf{p})) \quad \mathbf{u}(\tau) \in \mathbb{R}_+^n, \mathbf{p} \in \mathcal{D} \subseteq \mathbb{R}_+^\ell \\ \mathbf{u}(0) = \mathbf{u}_0 \in \mathbb{R}_+^n \end{cases} \tag{3}$$

where $\mathcal{D}$ is the parameter space and $\mathbf{u}(\tau, \cdot) = [P(\tau),\ I(\tau), A(\tau)]$ . Denote the associated time series data for each compartment as $\mathcal{P}_j, \mathcal{I}_j, \mathcal{A}_j$ Given $K$ observation times for host $j$ define

$$\mathbf{X_j} := [\mathcal{P}_{1,j},\ \ldots,\ \mathcal{P}_{K,j}, \mathcal{I}_{1,j},\ \ldots,\ \mathcal{I}_{K,j}, \mathcal{A}_{1,j},\ \ldots,\ \mathcal{A}_{K,j}] \in \mathbb{R}_+ \times \mathbb{R}_+^{3K} \tag{4}$$

We seek to find $\hat{\mathbf{p}} \in \mathcal{D}$ such that for

$$S_j(\mathbf{p}) = \sum_{s=1}^{3K} \left[ w_s(\mathbf{X}_j^{(s)} - \mathbf{u}_j^{(s)}(\tau, \mathbf{p})) \right]^2 \tag{5}$$

$$S_j(\mathbf{p}) \geq S_j(\hat{\mathbf{p}}) \tag{6}$$

for all $\mathbf{p} \in \mathcal{D}$.

In this formulation, the weights $\omega_s$ are treated as a hyperparameter manually chosen to reduce Average relative error while retaining the visual quality of fits. The host-specific values of $\omega_s$ are available in the program files archived in Zenodo (Macdonald et al., 2024).

To obtain these fit parameter point estimates, we numerically solve these two non-linear least squares problems in sequence using the interior-reflexive Newton method as implemented by MatLab's lsqcurvefit function. Beyond the measures of innate immunity shown in this work, we were unable to obtain quality fits for BKA data (see program files (Macdonald et al., 2024)) and use fits obtained from Haptoglobin as measures of innate response for comparative analysis across the route of infection, individual host variation, and sample mean parameter values and related quantities.

#### S4 Practical identifiability analysis and interval estimates

In order to assess identifiability, confidence in our fitting procedure, and differences in both mean *across and within serotype* we conducted uncertainty analysis via Monte Carlo simulations (see the methods section of the primary text), here, we outline how confidence intervals were obtained for serotype sample means.

1. For each host infected with a given serotype, we fit model (3), as described in the methods section of the primary text
2. Once a block of parameter estimates (4 per serotype, one from each host) has been generated, calculate the sample mean parameter values given those four drawn infection start times and the resulting parameter estimates
3. Arrange the generated values in increasing order and remove the top and bottom $2.5\%$ to obtain the desired approximate $95\%$ confidence intervals.

Inherent to this process is that the infection start times of each host are independent. While this is likely not strictly true, we cannot identify which host infected which host from a single infection experiment. Additionally, both serotype-specific transmission qualities *and* host-specific immunogenicity likely play a role in infection start time relative to the initiation of contact.

#### S5 Effect of route of infection on within-host viral and immune dynamics

Significant differences exist in viral and immune dynamic parameters between needle and contact-infected hosts within serotype (see supplementary table S3). In contrast, unlike the contact-infected hosts (table S1), there are few differences among serotypes for the needle-infected data (stable S2). Taken together, it indicates that needle-infected experiments alone cannot characterize the observed differences among serotypes in contact infection or a good characterization of individual serotype averages among contact-infected hosts.

Among all three serotypes, viral carrying capacity, the model parameter $K$, and maximum viral load (as calculated from time trajectory) were affected by the route of transmission, with SATs 1 and 2 displaying statistically significant differences [mean difference in carrying capacity for SAT1 -2.92, CI -4.39–1.29, for SAT2 -1.41, -2.63–0.27]. Particularly notable is the large difference in cumulative viral load for both SAT2 and SAT3 [mean difference -10.83, CI -14.46–6.95, 5.46, 0.91-10.16] respectively (See table S3 for all differences across routes of infection).

Differences in viral and immune dynamic parameters among serotypes are less detectable in needle infected hosts (see Supplementary Figures S1-S3). For needle infected hosts, differences among serotypes remain detectable for viral carrying capacity, $K$, maximum viral load, cumulative viral load, and adaptive activation and clearance rates. However these detected differences do not always line up with those detected in contact infected buffalo. Because of these differences in viral and immune dynamics between needle and contact infected hosts, we focus on contact infected hosts only for the remainder of this work.

#### S6 Supplementary Tables & Figures

Tables not included in the main text are provided below.

Table S1: Point estimates and confidence intervals for differences between sample means and sample means, *contact infected* hosts using *haptoglobin* as measure of innate immune response. Asterisk indicates difference between means is statistically significant

| Quantity | SAT1 – SAT2 | SAT1 – SAT3 | SAT2 – SAT3 |
|---|---|---|---|
| $k$ | 0.0387, (-0.107, 0.182) | 0.15, (0.00664, 0.281)* | 0.111, (-0.0325, 0.242) |
| $K$ | 3.07, (0.794, 5.06)* | -0.0174, (-2.45, 2.49) | -3.09, (-5.3, -0.572)* |
| $\delta$ | 0.0496, (-0.471, 0.545) | 0.132, (-0.419, 0.66) | 0.0827, (-0.513, 0.661) |
| $b$ | -0.118, (-0.18, -0.0668)* | -0.0615, (-0.162, -0.009)* | 0.0569, (-0.0531, 0.135) |
| $r$ | 0.736, (-0.919, 2.21) | 2, (0.53, 3.35)* | 1.27, (-0.333, 2.89) |
| $\nu$ | 2.36, (-0.623, 5.52) | -0.204, (-3.54, 3.26) | -2.56, (-5.27, 0.268) |
| $I_0$ | -0.298, (-1.24, 0.629) | -0.327, (-1.23, 0.531) | -0.029, (-0.952, 0.88) |
| $A_0$ | 0.139, (0.0812, 0.215)* | 0.14, (0.082, 0.216)* | 0.00127, (-0.0189, 0.0213) |
| $P_0$ | 0.00067, (-0.00142, 0.00254) | 0.00143, (-0.000681, 0.00316) | 0.000759, (-0.00111, 0.00249) |
| Max viral | 2.39, (0.586, 4)* | 0.365, (-1.59, 2.26) | -2.03, (-3.8, -0.102)* |
| Time max | -1.3, (-1.88, -0.609)* | -4.08, (-4.89, -3.18)* | -2.78, (-3.65, -1.89)* |
| Cum. Viral | 10.4, (3.82, 16.5)* | 1.69, (-11, 12.4) | -8.66, (-20.8, 1.57) |
| Start time | -0.675, (-1.47, 0.0828) | -1.6, (-3.51, -0.0381)* | -0.927, (-2.97, 0.84) |
| | SAT1 | SAT2 | SAT3 |
| $k$ | 0.3, (0.2, 0.404) | 0.261, (0.162, 0.365) | 0.15, (0.0773, 0.256) |
| $K$ | 8.39, (6.79, 9.89) | 5.32, (4.14, 7.02) | 8.41, (6.48, 10.2) |
| $\delta$ | 1.65, (1.34, 1.98) | 1.6, (1.22, 2.02) | 1.51, (1.11, 1.96) |
| $b$ | 0.102, (0.0852, 0.123) | 0.221, (0.174, 0.279) | 0.164, (0.116, 0.263) |
| $r$ | 4.44, (3.6, 5.46) | 3.71, (2.63, 5.12) | 2.44, (1.68, 3.63) |
| $\nu$ | 6.77, (4.28, 9.42) | 4.41, (2.81, 6.11) | 6.97, (4.74, 9.07) |
| $I_0$ | 3.31, (2.67, 3.92) | 3.6, (2.92, 4.28) | 3.63, (3.05, 4.22) |
| $A_0$ | 0.187, (0.132, 0.261) | 0.0479, (0.0336, 0.0627) | 0.0467, (0.033, 0.0619) |
| $P_0$ | 0.00788, (0.00632, 0.00926) | 0.00721, (0.006, 0.00869) | 0.00645, (0.00559, 0.00794) |
| Max viral | 7.19, (5.91, 8.41) | 4.8, (3.82, 6.16) | 6.83, (5.41, 8.27) |
| Time max | 3.14, (2.79, 3.62) | 4.44, (3.95, 4.88) | 7.22, (6.44, 7.95) |
| Cum. Viral | 31.2, (26.3, 35.9) | 20.8, (16.8, 25.3) | 29.5, (20.3, 40.8) |
| Start time | 0.376, (0.128, 0.661) | 1.05, (0.348, 1.81) | 1.98, (0.456, 3.85) |

Table S2: Confidence Intervals for differences between means and means, *needle infected hosts*. Asterisk indicates statistically significant difference.

| Quantity | SAT1 – SAT2 | SAT1 – SAT3 | SAT2 – SAT3 |
|---|---|---|---|
| $k$ | -0.0423, (-0.263, 0.181) | -0.0858, (-0.334, 0.155) | -0.0435, (-0.288, 0.187) |
| $K$ | 1.95, (-0.0797, 3.78) | 1.92, (-0.219, 3.88) | -0.0293, (-2, 2) |
| $\delta$ | 0.0361, (-0.333, 0.427) | -0.0169, (-0.346, 0.301) | -0.053, (-0.448, 0.3) |
| $b$ | -0.0414, (-0.0742, -0.0125)* | -0.0948, (-0.143, -0.0563)* | -0.0534, (-0.102, -0.00994)* |
| $r$ | -0.458, (-1.89, 0.984) | 0.382, (-0.652, 1.42) | 0.84, (-0.597, 2.28) |
| $\nu$ | 0.609, (-3.32, 4.56) | 1.29, (-2.4, 4.99) | 0.679, (-2.85, 4.27) |
| $I_0$ | 0.204, (-0.738, 1.11) | 0.142, (-0.804, 1.1) | -0.062, (-0.972, 0.851) |
| $A_0$ | 0.174, (0.122, 0.248)* | 0.16, (0.102, 0.236)* | -0.0143, (-0.0442, 0.0141) |
| $P_0$ | -3.86e-05, (-0.00109, 0.001) | 7.36e-05, (-0.000986, 0.00114) | 0.000112, (-0.000882, 0.00112) |
| Max viral | 0.754, (-0.965, 2.36) | 1.1, (-0.695, 2.76) | 0.341, (-1.36, 2.09) |
| Time max | 0.154, (-0.403, 0.665) | -0.00814, (-0.438, 0.417) | -0.162, (-0.66, 0.38) |
| Cum. Viral | -0.96, (-7.23, 5.23) | 9.71, (4.06, 15.2)* | 10.7, (5.03, 16.2)* |
| | SAT1 | SAT2 | SAT3 |
| $k$ | 0.381, (0.239, 0.559) | 0.423, (0.284, 0.583) | 0.466, (0.298, 0.662) |
| $K$ | 9.63, (8.08, 10.9) | 7.68, (6.45, 8.99) | 7.71, (6.19, 9.2) |
| $\delta$ | 2.05, (1.78, 2.25) | 2.01, (1.68, 2.25) | 2.06, (1.83, 2.25) |
| $b$ | 0.094, (0.0789, 0.115) | 0.135, (0.115, 0.164) | 0.189, (0.156, 0.234) |
| $r$ | 4.93, (4.24, 5.72) | 5.39, (4.19, 6.66) | 4.55, (3.91, 5.33) |
| $\nu$ | 7.46, (4.71, 10.4) | 6.85, (4.26, 9.63) | 6.17, (3.87, 8.7) |
| $I_0$ | 3.23, (2.55, 3.93) | 3.03, (2.42, 3.66) | 3.09, (2.44, 3.77) |
| $A_0$ | 0.235, (0.184, 0.308) | 0.0606, (0.0475, 0.0713) | 0.0749, (0.0499, 0.101) |
| $P_0$ | 0.00859, (0.00769, 0.00897) | 0.00862, (0.00777, 0.00895) | 0.00851, (0.00766, 0.00885) |
| Max viral | 7.95, (6.68, 9.01) | 7.2, (6.09, 8.37) | 6.85, (5.56, 8.12) |
| Time max | 2.45, (2.15, 2.76) | 2.3, (1.89, 2.75) | 2.46, (2.16, 2.76) |
| Cum. Viral | 30.5, (26.1, 34.8) | 31.5, (27, 36) | 20.8, (17.3, 24.1) |

Table S3: 95% confidence intervals for difference between sample means within serotype and across route of infection. The results indicate needle infected hosts are not useful for understanding contact transmission. Negative values indicate parameter value for needle hosts is greater, positive that contact infected value is greater.

| Quantity | SAT1 | SAT2 | SAT3 |
|---|---|---|---|
| $k$ | -0.0807, (-0.155, -0.0401)* | -0.162, (-0.219, -0.122)* | -0.316, (-0.407, -0.221)* |
| $K$ | -1.24, (-1.32, -1)* | -2.36, (-2.46, -1.97)* | 0.701, (0.286, 0.977)* |
| $\delta$ | -0.4, (-0.449, -0.274)* | -0.414, (-0.469, -0.235)* | -0.55, (-0.707, -0.287)* |
| $b$ | 0.00814, (0.00634, 0.00887)* | 0.0851, (0.0586, 0.115)* | -0.0252, (-0.0398, 0.0292) |
| $r$ | -0.49, (-0.641, -0.256)* | -1.68, (-1.73, -1.5)* | -2.11, (-2.27, -1.7)* |
| $\nu$ | -0.69, (-0.983, -0.386)* | -2.44, (-3.52, -1.43)* | 0.802, (0.362, 0.962)* |
| $I_0$ | 0.0738, (-0.00402, 0.118) | 0.576, (0.5, 0.623)* | 0.543, (0.448, 0.608)* |
| $A_0$ | -0.0478, (-0.0544, -0.0432)* | -0.0127, (-0.0146, -0.0085)* | -0.0283, (-0.0388, -0.017) |
| $P_0$ | -0.000706, (-0.00126, 0.000286) | -0.00141, (-0.00191, -0.000258)* | -0.00206, (-0.00248, -0.000904)* |
| Max viral | -0.758, (-0.807, -0.605)* | -2.4, (-2.47, -2.21)* | -0.0275, (-0.148, 0.152) |
| Time max | 0.688, (0.639, 0.859)* | 2.14, (2.06, 2.16)* | 4.76, (4.28, 5.2)* |
| Cum. Viral | 0.694, (0.217, 1.12)* | -10.6, (-10.8, -10.1)* | 8.71, (2.92, 16.7)* |

Table S4: Fixed host-specific innate response maintenance rates (model parameter $\Lambda$)

| Host ID | Group | Value | Host ID | Group | Value |
|---|---|---|---|---|---|
| 2 | SAT1 Contact | 0.1023 | 7 | SAT1 Needle | 0.1051 |
| 4 | SAT1 Contact | 0.1077 | 10 | SAT1 Needle | 0.1015 |
| 19 | SAT1 Contact | 0.0852 | 11 | SAT1 Needle | 0.1089 |
| 33 | SAT1 Contact | 0.1011 | 13 | SAT1 Needle | 0.1038 |
| 5 | SAT2 Contact | 0.1071 | 8 | SAT2 Needle | 0.0907 |
| 9 | SAT2 Contact | 0.096 | 20 | SAT2 Needle | 0.0847 |
| 22 | SAT2 Contact | 0.1048 | 28 | SAT2 Needle | 0.1046 |
| 29 | SAT2 Contact | 0.1055 | 32 | SAT2 Needle | 0.0916 |
| 12 | SAT3 Contact | 0.1084 | 26 | SAT3 Needle | 0.1018 |
| 15 | SAT3 Contact | 0.1036 | 27 | SAT3 Needle | 0.0978 |
| 16 | SAT3 Contact | 0.1094 | 34 | SAT3 Needle | 0.0717 |
| 17 | SAT3 Contact | 0.0971 | 35 | SAT3 Needle | 0.1047 |

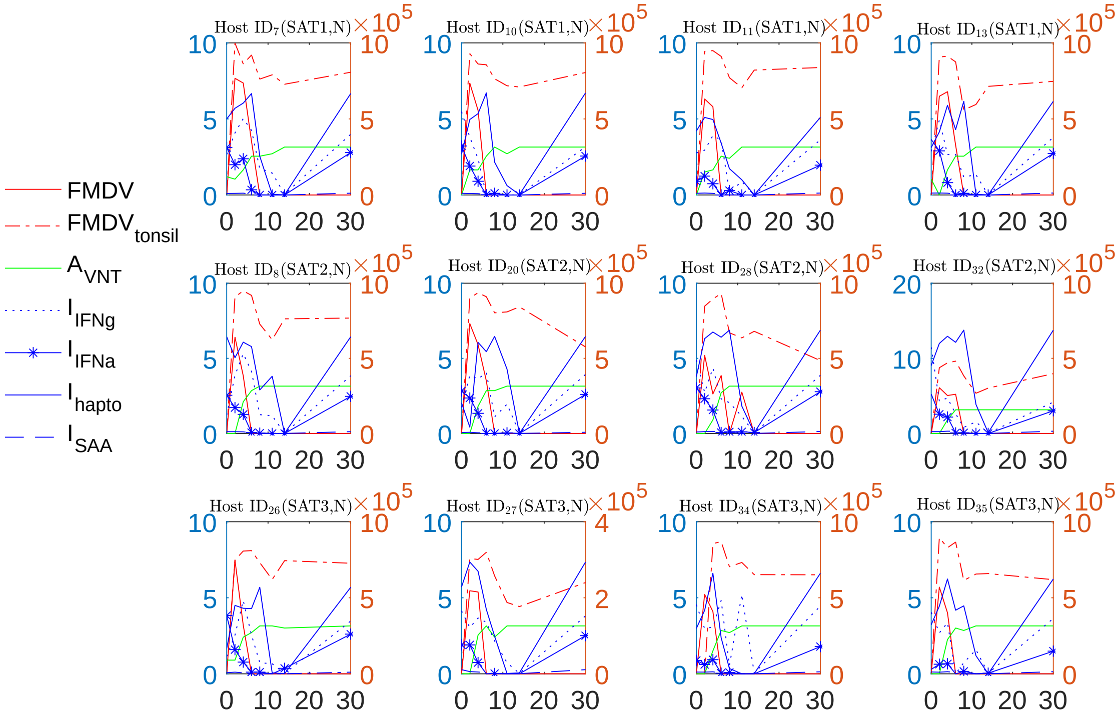

Figure S1: FMD virus and immune data, obtained from *needle infected hosts*. The right y-axis (orange axis) displays the concentration of innate immune responses $I_{hapto}$, and $I_{SAA}$. The rest of the virus-immune response concentrations is presented at the left y-axis (blue axis).

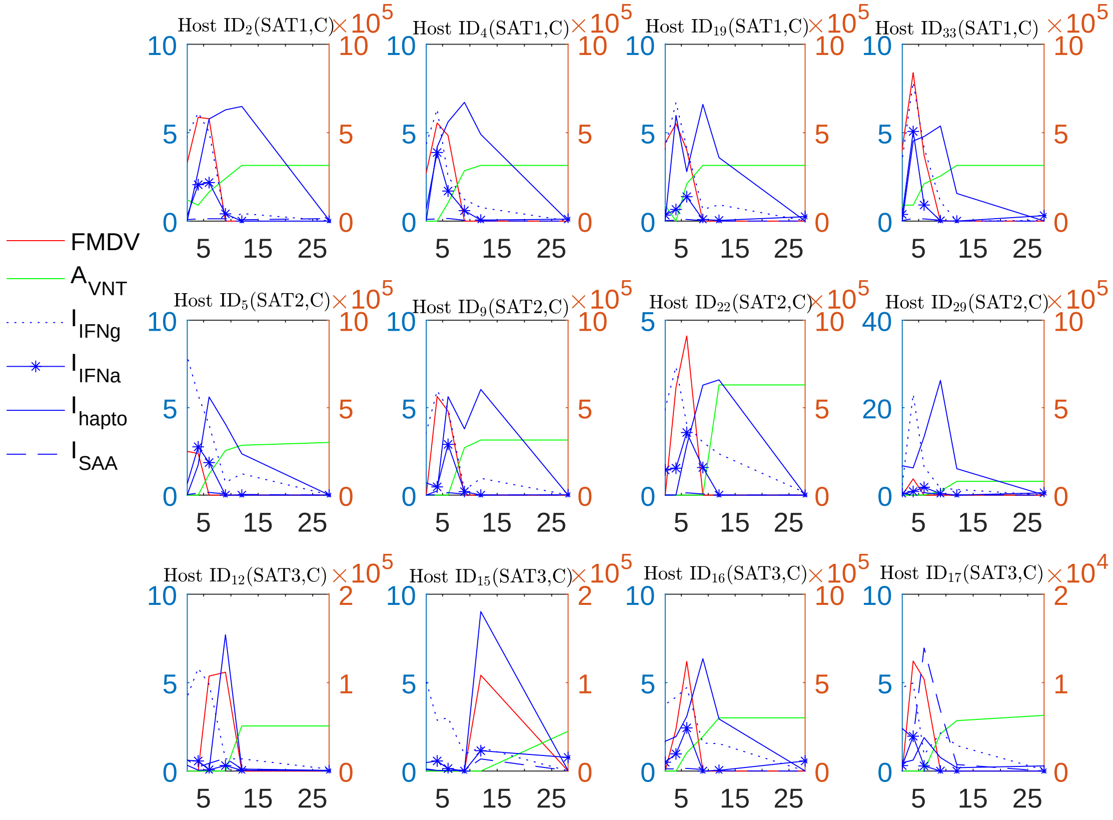

Figure S2: FMD virus and immune data obtained from *infected hosts via contact*. The right y-axis displays the concentration of innate immune responses $I_{hapto}$, and $I_{SAA}$. The rest of the virus-immune response concentrations is presented at the left y-axis.

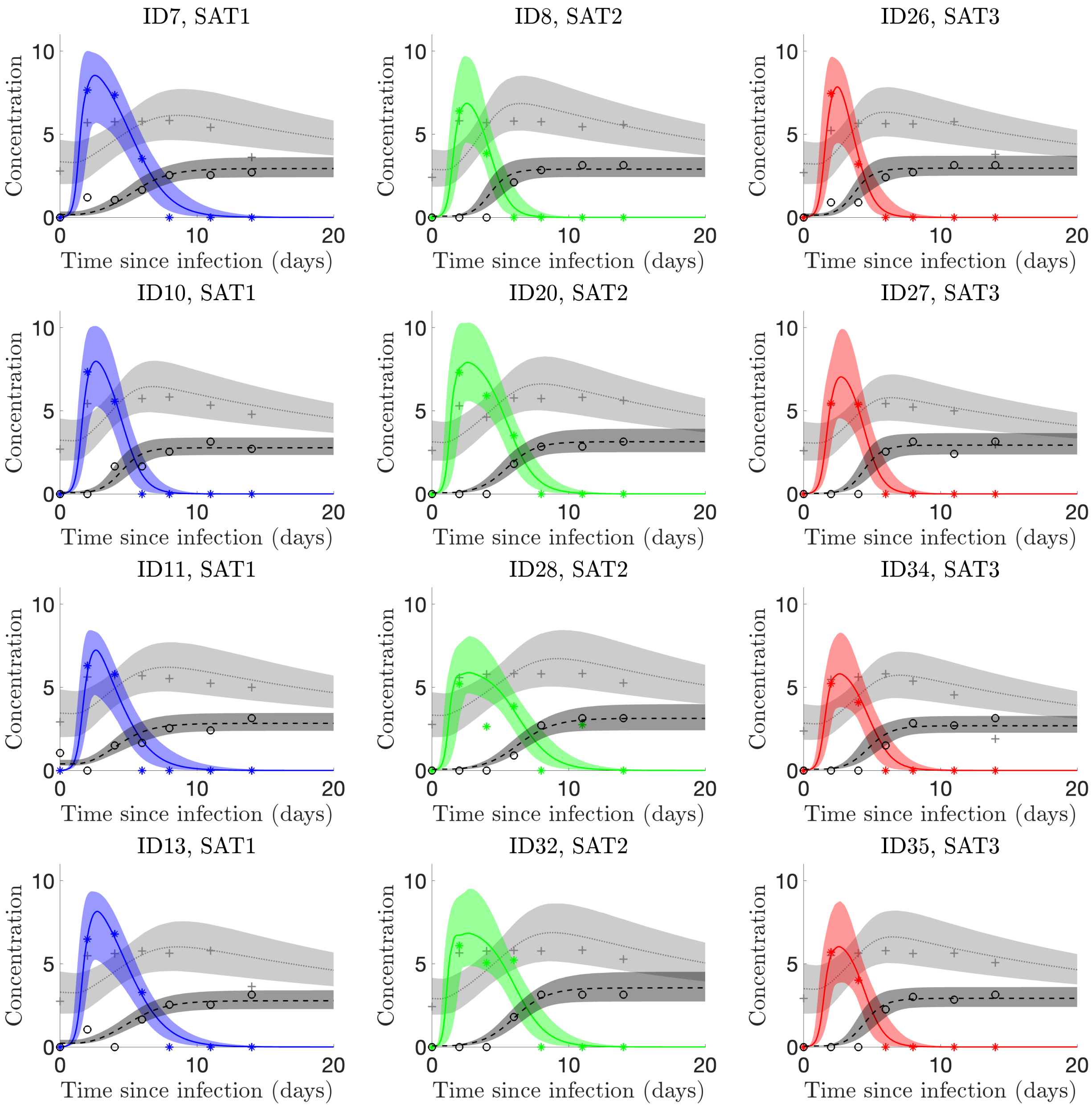

Figure S3: Our model reproduces empirically observed *needle infected* viral and immune dynamics well. Median time trajectories are represented by lines and 95% confidence intervals by shaded regions; blue indicates SAT1, green SAT2, and red SAT3. Haptoglobin is used to measure the innate response (light grey shade) [$\log_{10}(\mu g/mL)$], FMDV viral data is represented by colored shades [$\log_{10}$(genome copies/mL)], and virus neutralization titer [$\log_{10}$(VNT)] is used as a measure of adaptive response (dark grey shade). Virus data are indicated with asterisks, Haptoglobin with plusses, and VNT with circles.

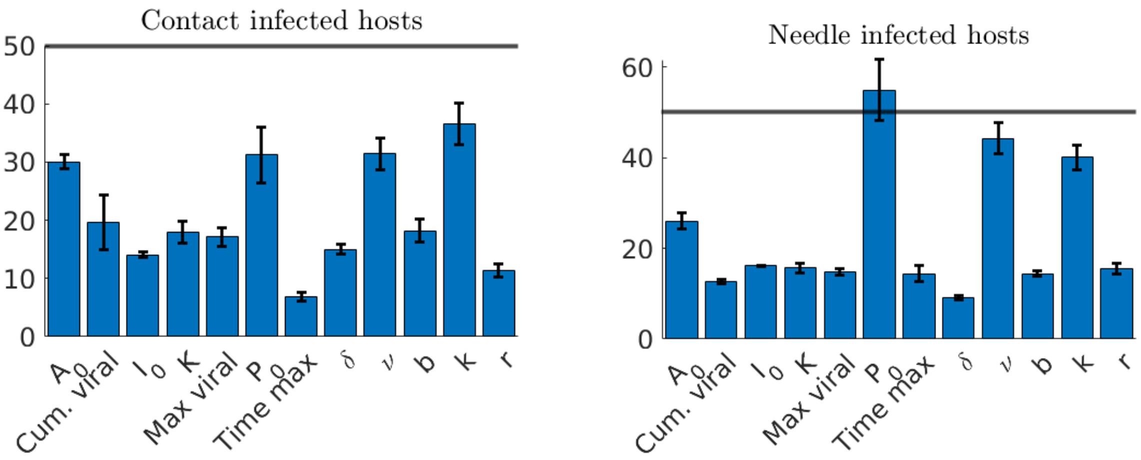

(a)

(b)

Figure S4: Identifiability analysis, (a) mean ARE for contact infected hosts with standard error, all model parameters are practically identifiable; (b) mean ARE for needle infected hosts with standard error, all parameters except initial viral load [$P_0$] are practically identifiable

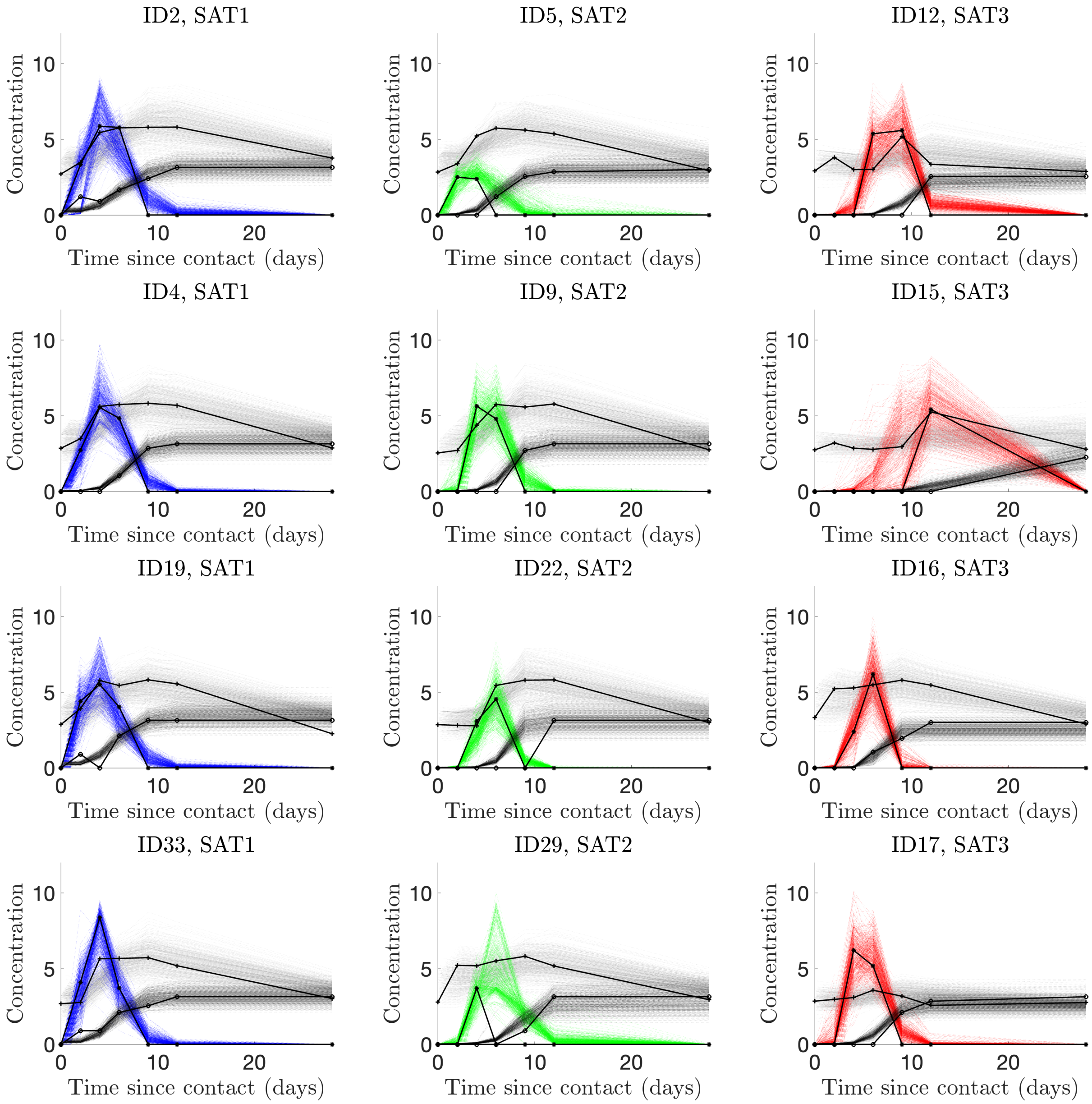

Figure S5: The choice of 50% noise covers the the empirical contact infection data barring clear outliers (that is $s = 0.5$ in equation (4) of the main text). The points connected with lines are the empirical data. The dotted lines are a simple random sample of 500 time solutions corresponding to fits to data generated by equation (4) in the main text. Virus data are indicated with asterisks, Haptoglobin with plusses, and VNT with circles.

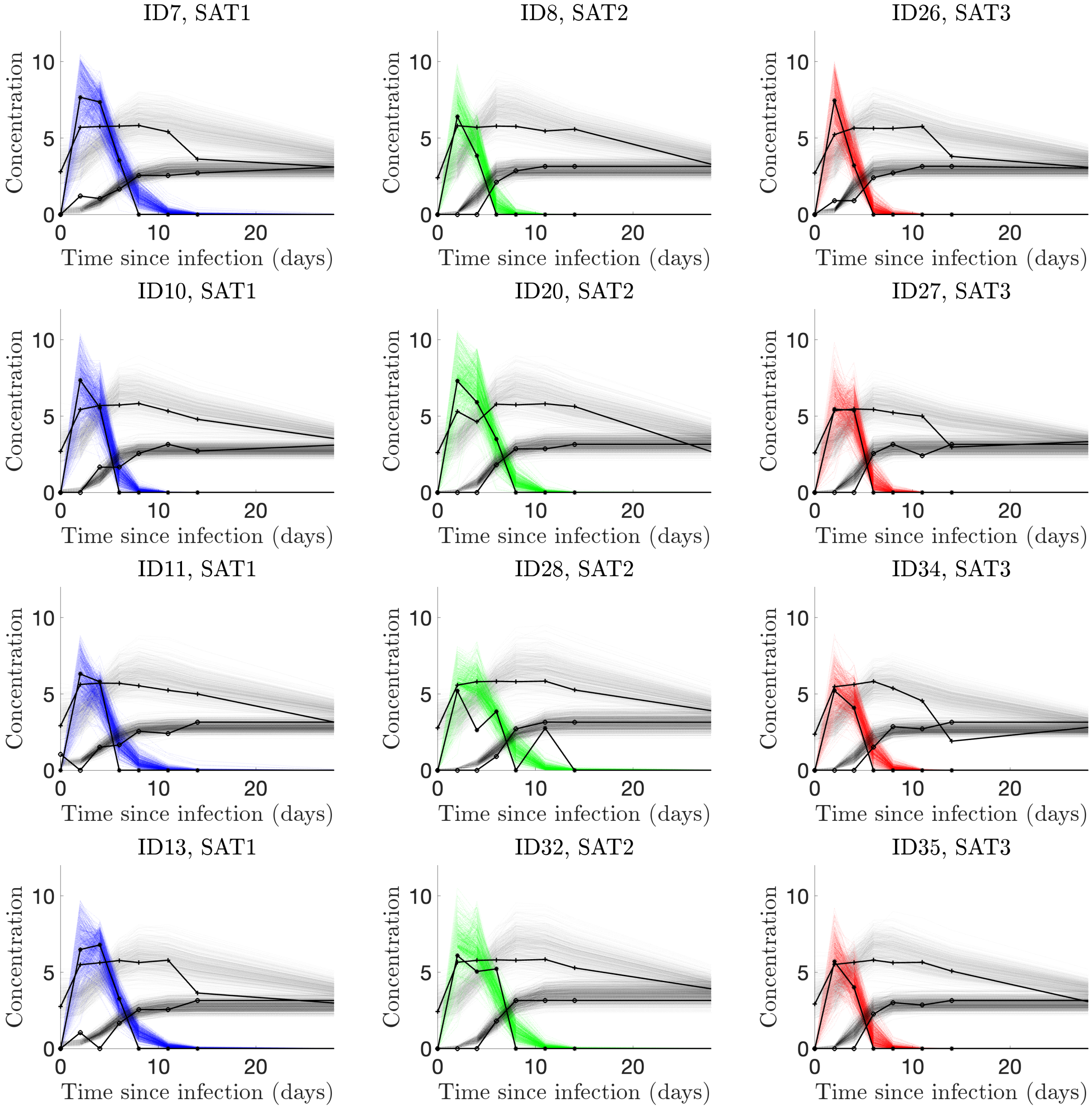

Figure S6: The choice of 50% noise covers the the empirical needle infection data barring clear outliers (that is $s = 0.5$ in equation (4) of the main text). The points connected with lines are the empirical data. The dotted lines are a simple random sample of 500 time solutions corresponding to fits to data generated by equation (4) in the main text. Virus data are indicated with asterisks, Haptoglobin with plusses, and VNT with circles.

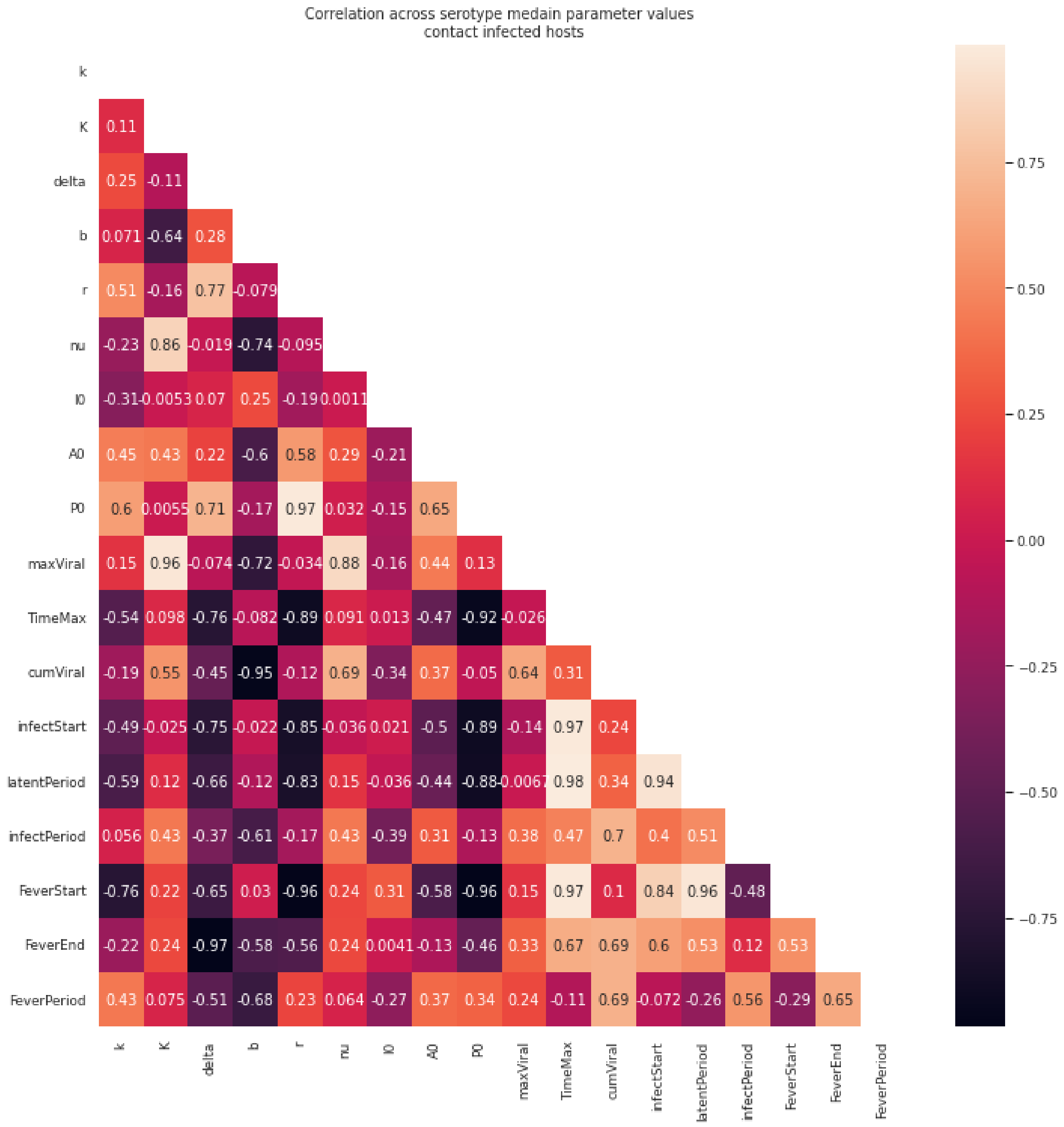

Figure S7: Pearson correlation between sample median values for 12 contact infected hosts
